# Mix-and-inject XFEL crystallography reveals gated conformational dynamics during enzyme catalysis

**DOI:** 10.1101/524751

**Authors:** Medhanjali Dasgupta, Dominik Budday, Saulo H.P. de Oliveira, Peter Madzelan, Darya Marchany-Rivera, Javier Seravalli, Brandon Hayes, Raymond G. Sierra, Sebastian Boutet, Mark Hunter, Roberto Alonso-Mori, Alexander Batyuk, Jennifer Wierman, Artem Lyubimov, Aaron S. Brewster, Nicholas K. Sauter, Gregory A. Applegate, Virendra K. Tiwari, David B. Berkowitz, Michael C. Thompson, Aina Cohen, James S. Fraser, Michael E. Wall, Henry van den Bedem, Mark A. Wilson

**Affiliations:** Department of Biochemistry and the Redox Biology Center, University of Nebraska, Lincoln, NE 68588, USA; Chair of Applied Dynamics, Friedrich-Alexander University Erlangen-Nürnberg, University of Erlangen-Nuremberg, D-91058 Erlangen, Germany; Bioengineering Department, Stanford University, Stanford, CA 94305, USA; Bioscience Division, SLAC National Accelerator Laboratory, Stanford University, Menlo Park, CA 94025, USA; Chemistry Department, University of Puerto Rico Mayaguez, Mayagüez PR 00681, USA; Linac Coherent Light Source, SLAC National Accelerator Laboratory, Stanford University, Menlo Park, CA 94025, USA; Stanford PULSE Institute, SLAC National Accelerator Laboratory, Stanford University, Menlo Park, CA 94025, USA; Stanford Synchrotron Radiation Lightsource, SLAC National Accelerator Laboratory, Stanford University, Menlo Park, CA 94025, USA; Molecular Biophysics and Integrated Bioimaging Division, Lawrence Berkeley National Laboratory, Berkeley, CA 94720, USA; Department of Chemistry, University of Nebraska, Lincoln, NE 68588, USA; Department of Bioengineering and Therapeutic Sciences, California Institute for Quantitative Biology, University of California, San Francisco, San Francisco, USA; Computer, Computational, and Statistical Sciences Division, Los Alamos National Laboratory, Los Alamos, NM 87505, USA

**Author notes:** **Author Contributions:** M.D. purified and crystallized ICH for synchrotron and XFEL experiments, assisted in synchrotron and XFEL data collection, processed synchrotron data, performed and analyzed enzyme kinetics experiments, drafted the manuscript; D.B. developed kinematic flexibility software and protocols, and carried out flexibility analysis; S.H.P.dO developed qFit software; P.M. purified and crystallized ICH; D.M.R performed the *in crystallo* UV-vis spectrophotometry and assisted in XFEL data collection; J.S. performed pre-steady state enzyme kinetics experiments and data analysis; B.H. assisted with *in crystallo* UV-vis spectrophotometry; R.G.S designed and operated the serial mixing sample delivery system at the LCLS; R.G.S. and A.B. helped prepare the experiment and MFX instrument; S.B., M.H., R.A.M. assisted with the XFEL experiment; J.W., A.L. processed the XFEL data; A.S.B. developed software for XFEL data processing and processed the XFEL data; N.K.S. developed software for XFEL data processing; G.A.A, V.K.T. and D.B.B. synthesized and validated the isocyanide substrate; M.C.T assisted in XFEL data collection; A.C. directed *in crystallo* UV-vis spectrophotometry and participated in XFEL data collection; J.S.F; directed XFEL data collection and drafted the manuscript; M.E.W. participated in XFEL experiment planning, MD simulations, and analysis; H.v.d.B. developed protein analysis software qFit and KGS, carried out molecular dynamics crystal simulations, and participated *in crystallo* UV-vis spectrophotometry. M.A.W. directed ICH purification and crystallization for synchrotron and XFEL data collection and directed enzyme kinetics data analysis. H.v.d.B. and M.A.W. conceived and designed the experiments and simulations, directed synchrotron and XFEL data collection, processing, analysis, and model refinement, analyzed all data, and drafted the manuscript.

**Keywords:** cysteine modification, radiation-controlled photo-oxidation, radiation damage, serial X-ray crystallography, qFit, enzyme conformational dynamics, serial mix-and-inject crystallography

## Abstract

**Summary Paragraph:** Protein dynamics play an important role in enzyme catalysis^1-4^. Many enzymes form covalent catalytic intermediates that can alter enzyme structure and conformational dynamics^5,6^. How these changes in enzyme structure and dynamics facilitate passage along the reaction coordinate is a fundamental unanswered question in structural enzymology. Here, we use <u>M</u>ix-and-Inject <u>S</u>erial Femtosecond X-ray <u>C</u>rystallography (MISC) at an X-ray Free Electron Laser (XFEL)^7-10^, ambient temperature X-ray crystallography, computer simulations, and enzyme kinetics to characterize how covalent modification of the active site cysteine residue in isocyanide hydratase (ICH) alters the enzyme’s conformational ensemble throughout the catalytic cycle. With MISC, we directly observe formation of a thioimidate covalent intermediate during ICH catalysis. The intermediate exhibits changes in the active site electrostatic environment, disrupting a hydrogen bond and triggering a cascade of conformational changes in ICH. X-ray-induced formation of a cysteine-sulfenic acid at the catalytic nucleophile (Cys101-SOH) with conventional crystallography at ambient temperature induces similar conformational shifts, demonstrating that these enzyme motions result from cysteine modification. Computer simulations show how cysteine modification-gated structural changes allosterically propagate through the ICH dimer. Mutations at Gly150 that modulate helical mobility reduce ICH catalytic turnover and alter its pre-steady state kinetic behavior, establishing that helical mobility is important for ICH catalytic efficiency. Taken together, our results demonstrate the potential of mix-and-inject XFEL crystallography to capture otherwise elusive mechanistic details of enzyme catalysis and dynamics from microcrystalline samples^7,11^. This approach can connect conformational dynamics to function for the large class of systems that rely on covalently modified cysteine residues for catalysis or regulation, resolving long-standing questions about enzyme mechanism and functionally relevant non-equilibrium enzyme motions.

---

The cysteine-dependent enzyme Isocyanide Hydratase (ICH: EC 4.2.1.103) is a homodimeric enzyme in the DJ-1 superfamily that hydrates diverse isocyanides to yield N-formamides^12^ (Figure 1a). Prior crystallographic studies of ICH showed evidence of helical mobility near the active site Cys101 residue (Figure 1A), which coincided with photooxidation of C101 to Cys101-SOH^13^. Cys101 makes an S^-^-HN hydrogen bond with the backbone amide of Ile152. Ile152 has a severely strained backbone conformation (φ =14°, ψ =-83°) that shifts to a relaxed, unstrained conformation of the displaced helix when this hydrogen bond is disrupted in a C101A mutant (Figure 1a)^13,14^.

**Figure 1:**
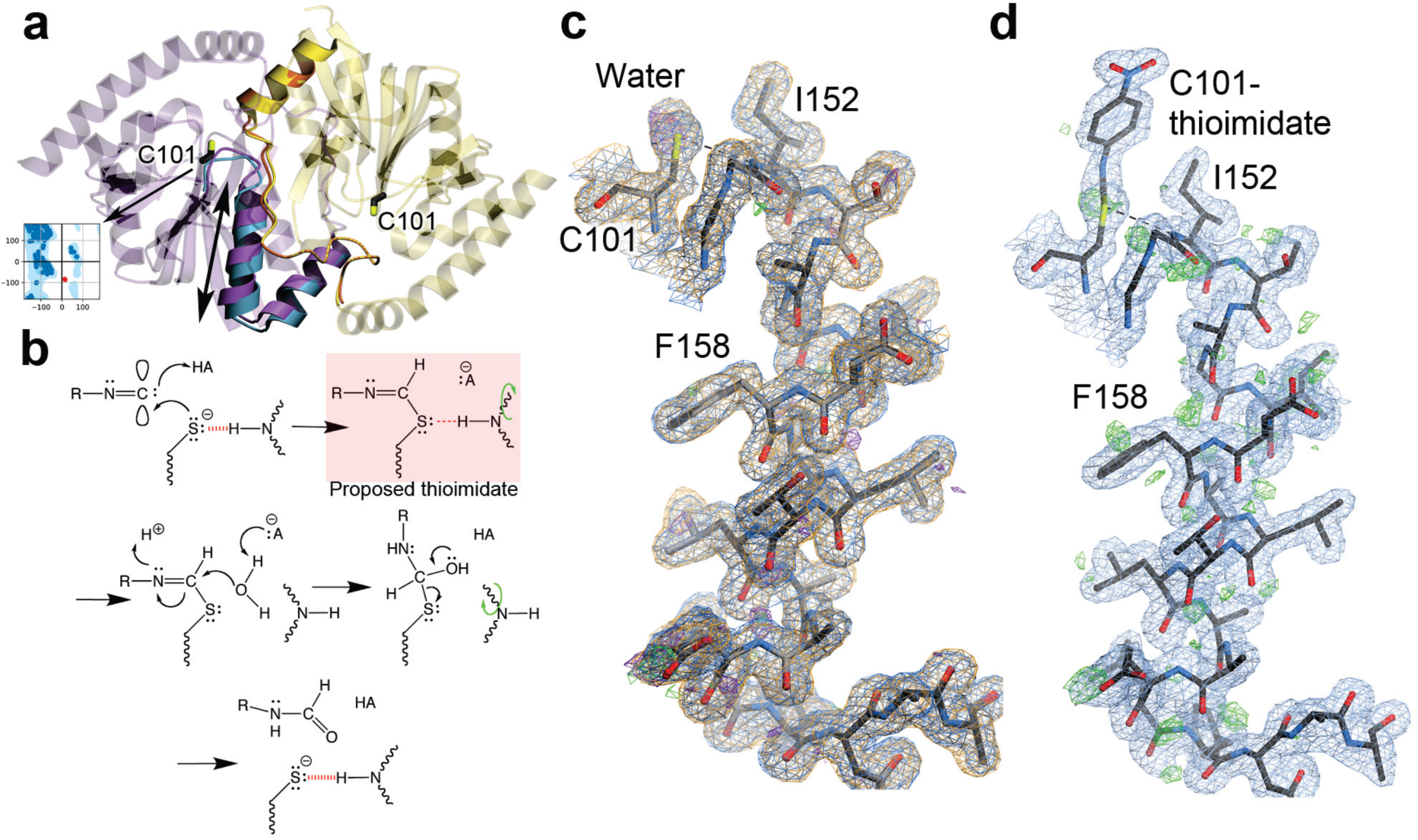
Catalytic intermediate captured with MISC. (a) The ICH dimer is shown as an overlay of WT ICH (purple) and C101A ICH (blue). The ‘B’ protomer is shown in yellow. The mobile helix H in C101A and areas exhibiting correlated backbone-sidechain disorder are rendered opaque. Ile152 is a Ramachandran outlier (inset) whose backbone torsion angles move with helical displacement. (b) Postulated reaction mechanism for ICH, beginning with Cys101 thiolate attack at the electrophilic carbenic carbon atom of isocyanide substrates and proceeding in the direction of the arrows.. A postulated thiomidate intermediate forms (red box), eliminating charge on Cys101 and weakening the H-bond to Ile152 (dashed red line). This relieves backbone torsional strain at Ile152 and permits sampling of shifted helix H conformations (green curved arrow), allowing water access to the intermediate for hydrolysis. (c) ICH completes a full catalytic cycle in the crystal during MISC. Helix H is shown with 2mF_o_-DF_c_ electron density (0.8 RMSD) prior to the introduction of substrate (blue) and after substrate has been exhausted (orange). These maps overlap almost perfectly, indicating ICH is fully restored to its resting conformation after catalysis. The hydrogen bond between the peptide backbone of Ile152 and Cys101 is shown in a dotted line. The helix is not mobile in these resting structures, indicated by the absence of features in the mF_o_-DF_c_ difference electron density (2.5 RMSD prior to substrate (green) and after catalysis is complete (purple) (d) ICH with a covalent thioimidate intermediate bound to Cys101 at the 15s time point. Difference mF_o_-DF_c_ electron density contoured at 2.5 RMSD (green) supports helix H sampling shifted conformations upon intermediate formation.

We examined the role of active site structural dynamics in ICH catalysis by directly monitoring catalysis in ICH microcrystals using MISC at 298K at carefully selected (Extended Data Figure S1) time delays at the LCLS XFEL, infusing the substrate p-nitrophenyl isocyanide (p-NPIC, Supplemental Information) into a stream of unmodified ICH microcrystals (Methods). The proposed reaction mechanism for ICH postulates that the catalytic Cys101 nucleophile attacks organic isocyanides at the electron deficient carbene-like center, followed by proton abstraction from a nearby general acid to generate a covalent thioimidate intermediate (Figure 1b). An enzyme-bound thioimidate, which has not been previously observed, eliminates the negative charge of the Cys101 thiolate (S^-^) and is proposed to reduce the strength of the Cys101-Ile152 hydrogen bond. Because SFX data are minimally affected by X-ray radiation damage^15^, the enzyme suffered no radiation-induced oxidation of Cys101 (Figure 1c). After 15 s of mixing, 2mF_o_-DF_c_ (Fig. 1d) and omit mF_o_-DF_c_, electron density maps (Extended Data Figure 2a) unambiguously reveal the formation of a thioimidate covalent intermediate in both ICH protomers. Once the intermediate is formed, positive mF_o_-DF_c_ difference electron density appears around helix H (Figure 1d) and its B-factors increase (Extended Data Figure 2b-d), indicating that it samples a new conformational ensemble. This difference density is absent both prior to catalysis and after ICH has exhausted substrate 5 minutes after mixing (Figure 1c). Therefore, helix H becomes transiently mobile upon intermediate formation during ICH catalysis.

**Figure 2:**
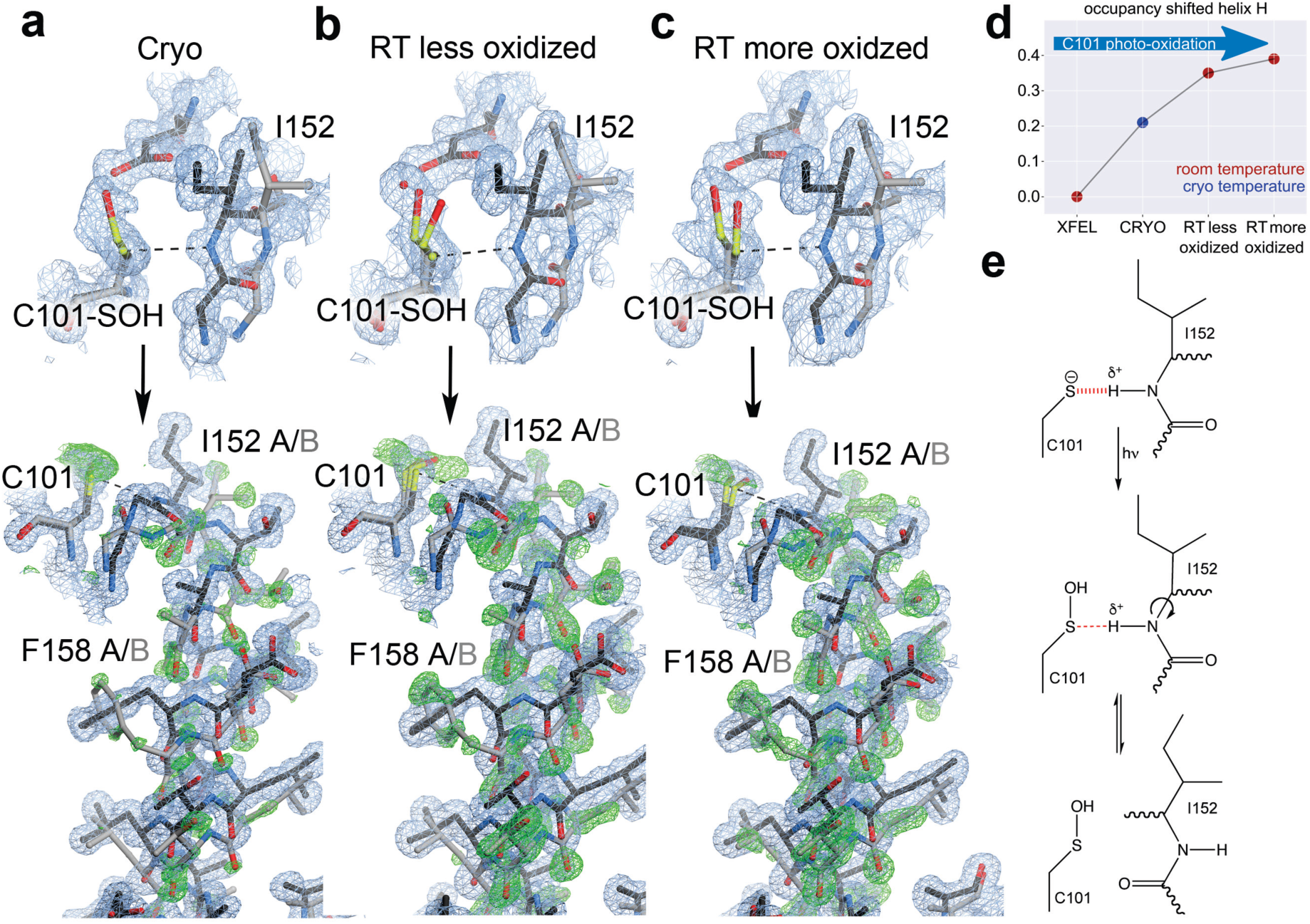
X-ray induced cysteine oxidation drives helical motion in ICH. (a-c) The top panels show the environment of Cys101 with varying degrees of oxidation to Cys101-sulfenic acid. 2mF_o_-DF_c_ electron density is contoured at 0.7 RMSD (blue) and the hydrogen bond between the peptide backbone of Ile152 and Cys101 is shown in a dotted line. “Cryo” is synchrotron data collected at 100 K (PDB 3NON), “RT less oxidized” is synchrotron data collected at 274 K with an absorbed dose of 2.4×10^4^ Gy, and “RT more oxidized” is synchrotron data collected at 277 K with an absorbed dose of 3.7×10^5^ Gy. The lower panels show helix H in its strained (black) and relaxed, shifted conformations (grey). 2mF_o_-DF_c_ electron density is contoured at 0.8 RMSD (blue) and omit mF_o_-DF_c_ electron density for the shifted helical conformation is contoured at 3.0 RMSD (green). At 274-277K, increased Cys101 oxidation disrupts the hydrogen bond to Ile152 and results in stronger difference electron density for the shifted helix conformation. (d) The refined occupancy of helix H for each X-ray dataset, indicating that increases in temperature and Cys101 oxidation result in higher occupancy for the shifted (relaxed) helix conformation. (e) Mechanism of X-ray induced covalent modification of C101 and weakening of the S^—^-NH hydrogen bond.

We reasoned that the on-pathway thioimidate must affect the interactions that hold helix H in a strained conformation similar to a covalent Cys101-SOH modification in the resting enzyme. Consistent with this idea, the cryogenic dataset collected at 100K (Cryo) also shows evidence of Cys101-SOH formation and a corresponding increase in helical mobility (Figure 2a)^13^. To enrich populations of the conformational shifts, we therefore collected a series of X-ray diffraction data sets of WT ICH at increasing doses of X-ray radiation (Figure 2a-c). Two room temperature (274-277K) synchrotron radiation datasets at 1.20-1.15 Å reveal a radiation-dose dependent Cys101 oxidation and helical displacement with increasing occupancy (Figure 2d). Notably, these stronger density peaks correspond to the difference electron density observed in the MISC experiment (Figure 2a-c). The origin of these shifts is radiation-induced oxidation of the Cys101, which eliminates the negative charge on S# thiolate and weakens the hydrogen bond between the amide H of Ile152 and Cys101 S#, from −2.2 kcal/mol with a thiolate acceptor to −0.91 kcal/mol with a Cys101-SOH acceptor (Figure 2e, Extended Data Figure S3; [Methods (3)]).

To investigate the local dynamical response of ICH to cysteine modification, we used molecular dynamics (MD) simulations of the reduced and Cys101-SOH crystals of ICH. We simulated the Cys101-SOH adduct rather than the thioimidate catalytic intermediate because it reports on the effects of charge neutralization (Figure 2e) more directly without the structural effects of the larger bound thioimidate intermediate or biases from thioimidate parameterization. Simulations of Cys101 thiolate (-S^-^) were started from the XFEL crystal structure (‘XFEL’ simulation, red) while the Cys101-SOH oxidized structure was similar to the synchrotron radiation-oxidized structure (‘SR’ simulation, blue). For the XFEL simulation, the Ile152-Cys101 thiolate hydrogen bond was predominantly maintained in the simulated crystal for the full 1 µs length of the MD trajectory (d_avg_(C101_SG_,I152_H_) = 2.9Å, Figure 3a). By contrast, in simulations of the SR Cys101-SOH state the hydrogen bond typically dissociated very early in the trajectory (d_avg_(C101_SG_,I152_H_) = 4.0Å, Figure 3A). Moreover, several protomers in the simulated Cys101-SOH crystal experienced a shift of helix H similar to that observed in the ambient temperature synchrotron radiation datasets (mean shift 0.69Å, Figure 3b, movie S1). These helical shifts were observed less frequently in simulations with the Cys101 thiolate (mean shift 0.60Å, Figure 3b), consistent with the hypothesis that local redistribution of cysteine electrostatic charge modulates these long-range motions.

**Figure 3:**
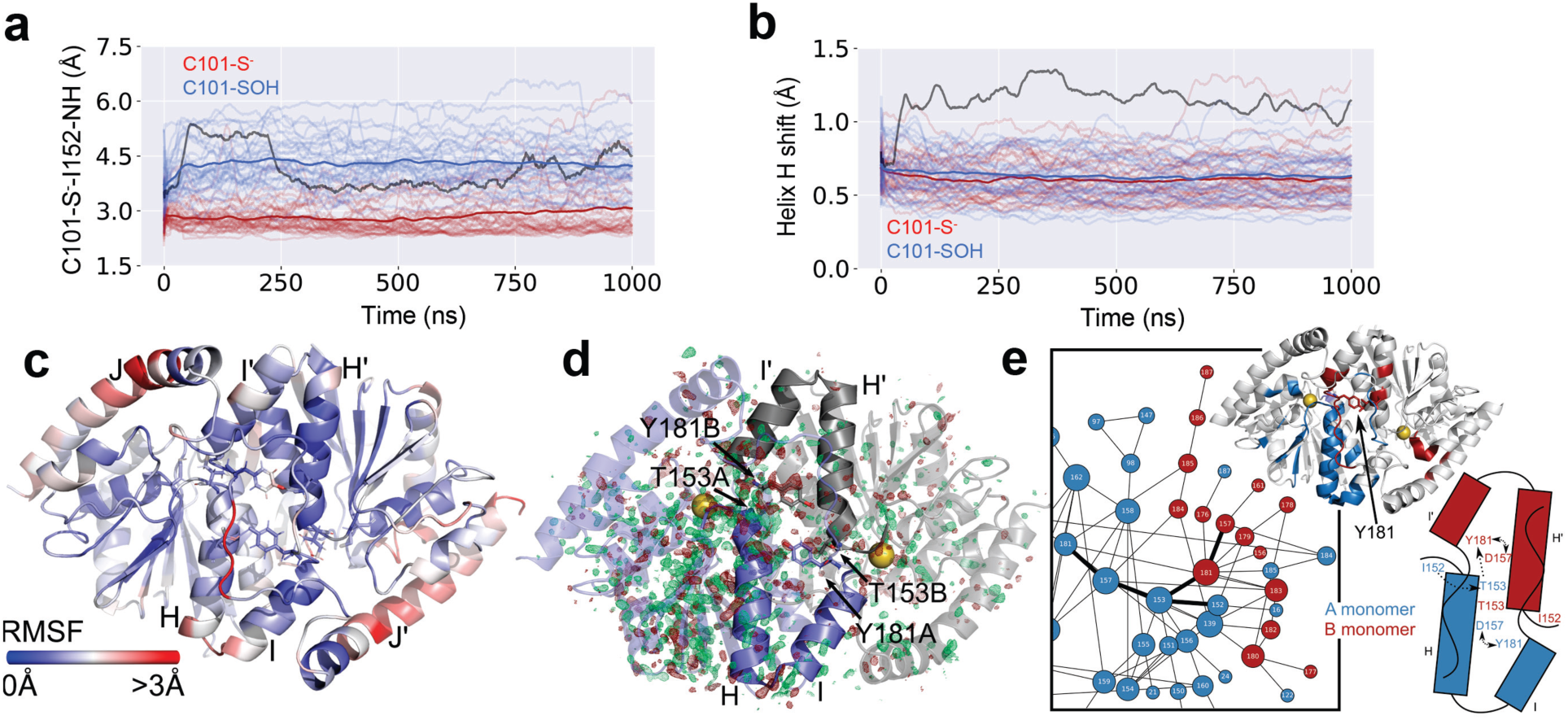
Cysteine modification results in a protein-wide conformational response. (a) Fluctuations of the C101_SG_-I152_H_ distance in simulations of ICH crystals with Cys101 in the thiolate (red) or Cys101-SOH (blue) state. Each ICH dimer is represented by a transparent line. Opaque red and blue lines denote the average C101_SG_-I152_H_ distance across the dimers. (b) Conformational shift of helix H with Cys101 in the thiolate (red) or Cys101-SOH (blue) state. The dark grey lines in panels (a) and (b) represent the trajectory selected for Extended Data Movie S1. (c) Root Mean Square Fluctuations calculated from MD simulations indicate highest fluctuations in linker I-J of the B protomer. Helix H of the A protomer just underneath the linker also shows elevated RMSF. (d) An isomorphous F_o_(SR) – F_o_(XFEL_APO_) difference map reveals features (green, positive; red, negative) distributed throughout the dimer, suggesting broadly altered structure and dynamics upon formation of the cysteine-sulfenic acid in the 274K synchrotron radiation (F_oRT_) dataset. The ‘A’ conformer is shown in slate, and the ‘B’ conformer in grey transparent cartoon representation. Helices H and I are shown opaque in both conformers. The catalytic nucleophile is shown in spheres. Difference electron density features are non-uniformly distributed, with stronger features near helix H in the A conformer, and along region B169-B189, which contacts the N-terminal end of helix H. Isomorphous difference features were calculated using an F_o_ sigma cutoff of 2.0 in both cases and maps were contoured at +/-3.0 RMSD. (e) CONTACT analysis identifies allosteric coupling across the dimer interface, in striking agreement with isomorphous difference maps and RMSF from panel (d). The A protomer is color-coded in blue, the B protomer in red. Residues identified in the CONTACT analysis are projected onto the cartoon representation.

Root Mean Square Fluctuations (RMSFs) calculated along a simulation trajectory show wide spread movement across the entire dimer (Figure 3c). Consistent with this simulation result, an F_o_(SR)–F_o_(XFEL_APO_) isomorphous difference electron density map of the Cys101-SOH SR dataset (F_o_ (SR)) and the Cys101-S^-^ XFEL dataset before substrate was introduced (F_o_(XFEL_APO_)) revealed widely distributed difference features (Figure 3d). An F_o_(XFEL_15s_)– F_o_(XFEL_APO_) difference map revealed similar, but weaker features suggesting that formation of the intermediate at 15s after mixing similarly shifts the conformational ensemble (Extended Data Figure S4). The F_o_(SR) – F_o_(XFEL_APO_) difference map features are most prominent in protomer A near the active site and the mobile helix H (Figure 3d). Difference features at the active site correspond to the dramatic conformational change of a conserved diglycyl motif G150-G151 and Ile152, while those near the interface of helix H and the $-sheet in protomer A reflect residues adjusting their position in response to helix H motion. There are far fewer peaks in protomer B, consistent with the absence of mobility of helix H’ in that protomer. However, the F_o_(SR)–F_o_(XFEL_APO_) map reveals significant features along the C-terminus of helix I’ at the dimer interface in protomer B, which directly contacts ThrA153 through TyrB181 (Figure 3d). These peaks likely report on conformational changes allosterically propagating from protomer A through the I’J’ linker (residues 181-185) into protomer B. By contrast, the difference map around the C-terminus of helix I and linker IJ in protomer A (IJ_A_, which packs against the stationary helix H of protomer B) is comparatively featureless.

Interestingly, the simulations indicate that the conformational shift in linker I’J’ towards the unshifted helix H’ and corresponding active site is larger than observed in the crystal structure (Extended Data Movie S1). Notably, our simulations suggest that the I’J’ conformational shift can precede relaxation of the strained Ile152 conformation and subsequent shift of helix H by several nanoseconds, allosterically communicating dynamical changes in protomer A across the dimer interface into protomer B. We further examined the dynamical communication across the dimer interface in ICH using CONTACT network analysis. CONTACT elucidates pathways of collective amino acid main- and sidechain displacements through mapping van der Waals conflicts in multiconformer qFit models^16^ that would result from sidechain conformational disorder if correlated motions are not considered^17^. CONTACT identified a large network of correlated residues in protomer A (with the mobile helix) that connects with a smaller network in protomer B (Figure 3e), corroborating the isomorphous difference map. We then used Kinematic Flexibility Analysis (KFA,^18^) to analyze how the Cys101-Ile152 hydrogen bond modulates motion modes accessible to the enzyme (Extended Data Figure S5)). We compared motion modes corresponding to the lowest mode-specific free energies in the XFEL structure when the H-bond is intact in both A and B protomers (C101-I152_A&B_) to when this H-bond is disrupted in protomer A (C101-I152_B_) (Extended Data Figure S6)). These altered motion modes are those most affected by disruption of the hydrogen bond. Perturbations in the hydrogen bonding network are propagated primarily to the IJ-linkers, helix H and I and helices J and J’, consistent with the MD simulations. Strikingly, KFA predicted large RMSFs within the IJ_A_ linker near the active site in protomer B, suggesting that the two sites are in allosteric communication. Considered together, the isomorphous difference electron density map, CONTACT analysis, KFA, and long-time MD simulations all point to local changes in hydrogen bonding at Cys101 initiating a cascade of conformational changes that propagate across the entire ICH dimer in an asymmetric manner^4^.

To test how helical mobility is related to ICH catalysis, we designed two mutations, G150A and G150T. G150 is part of a highly conserved diglycyl motif that moves ∼3Å to accommodate helical motion in ICH. Ambient temperature (274-277 K) crystal structures of G150A and G150T ICH show that the added steric bulk at this position biases the helix towards its relaxed conformation (Figure 4). In G150A ICH, helix H samples both conformations in both protomers (Figure 4a, Extended Data Figure S7). By contrast, the G150T structure shows helix H constitutively shifted to its relaxed position (Figure 4b). Both G150A and G150T have an alternate Cys101 sidechain conformation which conflicts with the unshifted conformation of Ile152 and helix H (Figure 4a,b, asterisk), indicating that the helix must move partially independently of Cys101 oxidation in these mutants. As in wild-type ICH, G150A shows evidence of Cys101-SOH oxidation in the electron density (Figure 4a). In contrast, Cys101 in G150T ICH is not modified by comparable exposure to X-rays (Figure 4b), indicating that the hydrogen bond donated to Cys101-S^-^ is important for enhancing Cys101 reactivity.

**Figure 4:**
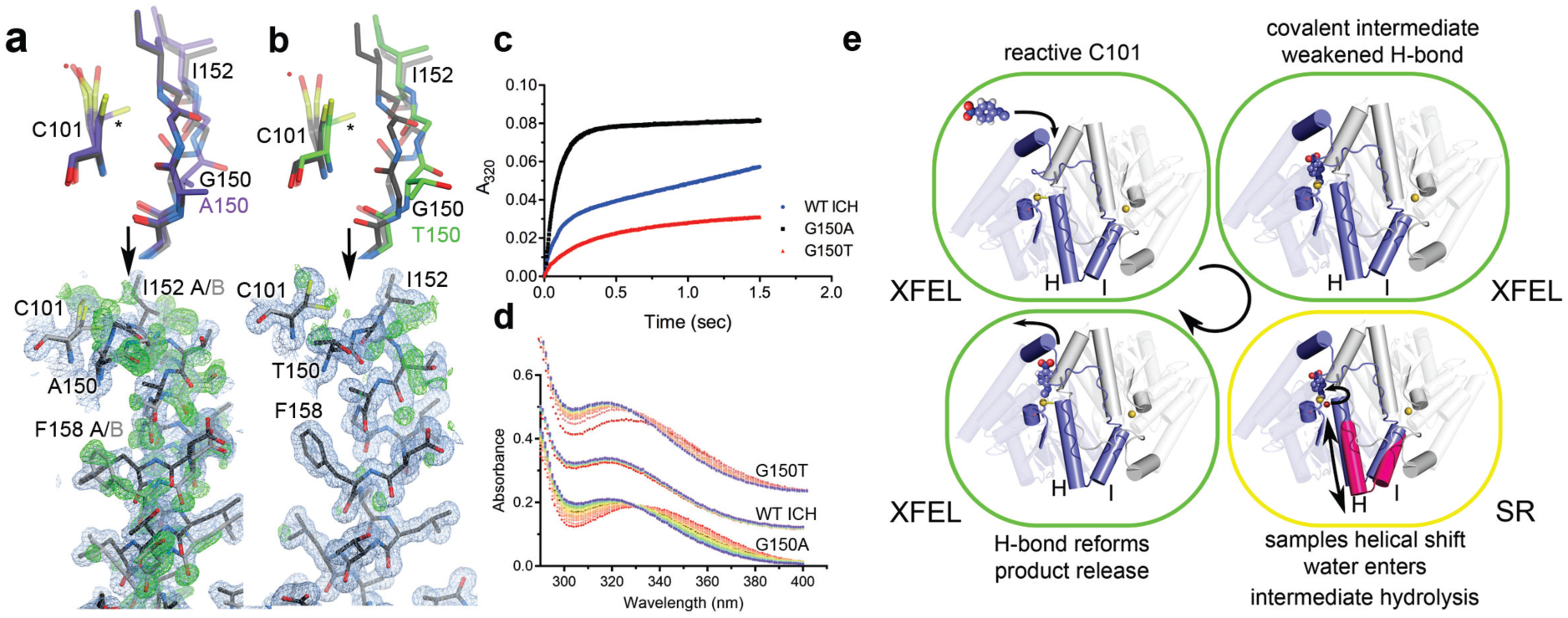
Mutations at Glu150 alter helical mobility and reduce ICH catalytic turnover. (a, b): The top portions show the environment of Cys101 in Gly150A and G150T ICH. 2mF_o_-DF_c_ electron density is contoured at 0.7 RMSD (blue) and the hydrogen bond between the peptide backbone of Ile152 and Cys101 is shown in a dotted line. Both G150 mutations permit unmodified Cys101 to sample conformations (asterisk) that sterically conflict with Ile152 in the strained helical conformation (black). Therefore, the helix in G150A and G150T ICH must sample shifted conformations (grey) in the absence of Cys101 modification. The lower portions of (A) and (B) show the helix in its strained (black) and relaxed, shifted conformations (grey). 2mF_o_-DF_c_ electron density is contoured at 0.8 RMSD (blue) and omit mF_o_-DF_c_ electron density for the shifted helical conformation is contoured at +3.0 RMSD (green). The helix samples both strained and shifted conformations in G150A ICH, while the helix is constitutively shifted in G150T. Pre-steady state (c) enzyme kinetics of wild-type (WT; blue circles), G105A (black squares), and G150T (red triangles) ICH. Pre-steady kinetics at 160 μM p-NPIC show a pronounced burst phase for each protein, but with variable burst and steady state rate constants (k_burst_(WT)= 11.395±0.120 s^-1^, k_burst_(G150A)= 11.882±0.071 s^-1^, k_burst_(G150T)= 4.255±0.187 s^-1^). The divergent pre-steady state profiles indicate that G150A impacts steps after the first chemical step, while G150T affects both early and later steps. (e) Single turnover spectra of ICH enzymes with p-NPIC substrate shown as a function of time, from early (red) to later (blue) timepoints. Spectra were collected every five seconds. At early times, G105A and G150T accumulate a species with λ_max_=335 nm, likely the thioimidate intermediate that resolves to product in the blue spectra with λ_max_=320 nm. (e) Clockwise from upper left: substrate enters the protomer with the active site Cys101 thiolate poised for nucleophilic attack and the Cys101-Ile152 H-bond intact. Formation of the thioimidate intermediate weakens the Cys101-Ile152 H-bond and causes helical motion (magenta helices H and I) that promotes intermediate hydrolysis (red sphere; curved arrow). Hydrolysis of the thioimidate intermediate restores the reactive C101 thiolate and strengthens the Cys101-Ile152 H-bond, thereby shifting the helix conformational ensemble to favor the strained conformation.

Unlike the C101A mutant, which shows similar helical displacement but is catalytically inactive, G150A and G150T ICH are catalytically active. This allowed us to investigate the role of helical displacement in the ICH catalytic cycle. Steady-state enzyme kinetics of the G150A and G150T mutants measured using p-NPIC as the substrate show a ∼6–fold reduction in k_cat_ for both mutants compared to wild-type enzyme but largely unchanged K_M_ values (Extended Data Fig. S8). In contrast to their similar steady-state kinetic behavior, the G150 mutants have divergent pre-steady state kinetic profiles in stopped-flow mixing. ICH exhibits “burst” kinetics (Figure 4c), indicating that the rate-limiting step for ICH catalysis comes after formation of the thioimidate intermediate. G150A ICH has a burst exponential rate constant k of ∼11 s^-1^ that is comparable to the wild-type enzyme, but G150T has a reduced burst rate constant of ∼4 s^-1^, indicating a slower chemical step (Figure 4c, Extended Data Figure S8). WT and G150A ICH have comparable second order rate constants for the burst phase, while G150T is markedly slower (Extended Data Figure S8). Therefore, although both the G150A and G150T mutations impair ICH catalysis, the kinetic effect of the G150A mutation is predominantly in steps after formation of the intermediate, while G150T impairs both the rate of intermediate formation and later, rate-limiting steps. Furthermore, we observe spectral evidence for the thioimidate intermediate in single-turnover UV-visible spectra (Figure 4D). G150A ICH accumulates a species whose absorbance maximum is 335 nm, while WT ICH completes a single turnover and accumulates the 320 nm formamide product in the ∼30 second deadtime of manual mixing (Figure 4D). G150T ICH accumulates less of the 335 nm intermediate than G150A ICH due to a closer match between the rates of formation and consumption of the intermediate in G150T ICH. In G150A ICH, the 335 nm species slowly converts to the 320 nm product over ∼40 s (Figure 4D), consistent with the slow rate of product formation observed after the burst in G150A pre-steady state kinetics (Figure 4c). These data support the conclusion that the 335 nm species is the ICH-thioimidate intermediate observed in the MISC experiment and that G150A is impaired in hydrolyzing this covalent intermediate from the active site Cys101.

Covalent modification is a common and physiologically important perturbation to proteins. Reactive residues such as cysteine are prone to diverse covalent modifications with catalytic or regulatory consequences. We found that cysteine modification in ICH remodels active site H-bonding networks and gates catalytically important changes in protein dynamics. Modification of the active site cysteine thiolate during catalysis neutralizes its negative charge and weakens a key H-bond, initiating a cascade of conformational changes in ICH that span the entire dimer. Transiently increased mobility in the active site facilitates water entry and attack at the thioimidate to form a proposed tetrahedral intermediate. The H-bond between Cys101 and Ile152 then reforms, stabilizing the nascent Cys101 thiolate and making it a better leaving group, thereby coupling dynamical changes in the enzyme to progress along the reaction coordinate (Figure 4e). Notably, all cysteine thiolates will experience a similar loss of negative charge upon covalent bond formation. Because cysteine residues can have multiple roles in a protein, including catalytic nucleophile, metal ligand, acylation target, redox target, and others, many different modification-mediated signals may be transduced through altered cysteine electrostatics to impact protein dynamics and function, expanding the ways in which cysteine can couple protein biophysical properties to cellular needs. Integrating recent developments in serial and conventional RT crystallography with advanced computational methods^19^ can offer new, exciting opportunities to reveal enzyme mechanism and interrogate the consequences of cysteine modification on protein dynamics.

## Acknowledgements

We thank Dr. Donald Becker (University of Nebraska) and Dr. Joseph Barycki (North Carolina State University) for helpful discussions and Lauren Barbee (University of Nebraska) for assistance with ICH kinetic data and analysis. RGS acknowledges the support of the OBES through the AMOS program within the CSGB and of the DOE through the SLAC Laboratory Directed Research and Development Program. ASB and NKS were supported by NIH grant GM117126 to NKS for data-processing methods. MCT and JSF were supported by NSF (STC-1231306), by a Ruth L. Kirschstein National Research Service Award (F32 HL129989) to MCT, and by awards to JSF from NIH (GM123159, GM124149), the David and Lucile Packard Foundation (Packard Fellowship), and the UC Office of the President Laboratory Fees Research Program (LFR-17-476732. MEW was supported by the Exascale Computing Project (17-SC-20-SC), a collaborative effort of the US Department of Energy Office of Science and the National Nuclear Security Administration. SHPdO and HvdB are supported by award NIH GM123159 to HvdB. HvdB is supported by a Mercator Fellowship from the Deutsche Forschungsgemeinschaft (DFG). MAW was supported by Nebraska Tobacco Settlement Biomedical Research Development Fund. Use of the Stanford Synchrotron Radiation Lightsource, SLAC National Accelerator Laboratory, is supported by the U.S. Department of Energy, Office of Science, Office of Basic Energy Sciences under Contract No. DE-AC02-76SF00515. The SSRL Structural Molecular Biology Program is supported by the DOE Office of Biological and Environmental Research, and by the National Institutes of Health, National Institute of General Medical Sciences (including P41GM103393). Use of the Linac Coherent Light Source (LCLS), SLAC National Accelerator Laboratory, is supported by the U.S. Department of Energy, Office of Science, Office of Basic Energy Sciences under Contract No. DE-AC02-76SF00515. This research used resources of the Advanced Photon Source, a U.S. Department of Energy (DOE) Office of Science User Facility operated for the DOE Office of Science by Argonne National Laboratory under Contract No. DE-AC02-06CH11357. Use of BioCARS was also supported by the National Institute of General Medical Sciences of the National Institutes of Health under grant number R24GM111072. The contents of this publication are solely the responsibility of the authors and do not necessarily represent the official views of NIGMS or NIH.

## Methods

### Crystallization, data collection, and processing

*Pseudomonas fluorescens* ICH was expressed in *E. coli* as a thrombin-cleavable, N-terminally 6xHis-tagged protein and purified as previously described^13^. All of the final proteins contain the vector-derived amino acids “GSH” at the N-terminus. Wild-type ICH, the G150A, and the G150T mutants were crystallized by hanging drop vapor equilibration by mixing 2 µL of protein at 20 mg/ml and 2 µl of reservoir (23% PEG 3350, 100mM Tris-HCl, pH 8.6, 200 mM magnesium chloride and 2 mM dithiotheritol (DTT)) and incubating at 22°C. Spontaneous crystals in space group P2_1_ appear in 24-48 hours and were used to microseed other drops after 24 hours of equilibration. Microseeding was used because two different crystal forms of ICH crystals grow in the same drop and those in space group P2_1_ are the better-diffracting crystal form. Notably, seeding G150T ICH using wild-type ICH crystals in space group P2_1_ results exclusively in G150T crystals in space group C2 with one molecule in the asymmetric unit (ASU).

Cryogenic (100K) synchrotron data (PDB 3NON) were collected at the Advanced Photon Source beamline 14BM-C from plate-shaped crystals measuring ∼500×500×150 µm that were cryoprotected in 30% ethylene glycol, mounted in nylon loops, and cooled by immersion in liquid nitrogen as previously described^13^. Ambient temperature (274 K and 277 K) synchrotron data sets were collected at the Stanford Synchrotron Radiation Lightsource from crystals mounted in 0.7 mm diameter, 10 µm wall thickness glass number 50 capillaries (Hampton Research) with a small volume (∼5 µl) of the reservoir to maintain vapor equilibrium. Excess liquid was removed from the crystal by wicking, and the capillary was sealed with beeswax. For the 274 K dataset, the crystal was mounted with its shortest dimension roughly parallel to the capillary axis, resulting in the X-ray beam shooting through the longest dimensions of the crystal during rotation. For the 277 K dataset, a large single capillary-mounted crystal was exposed to X-rays and then translated so that multiple fresh volumes of the crystal were irradiated during data collection. This strategy reduces radiation damage by distributing the dose over a larger volume of the crystal. This translation of the sample was factored into the absorbed dose calculation by assuming every fresh volume of the crystal received no prior dose, which is a best-case scenario.

The 100 K and 277 K datasets were collected on ADSC Q4 CCD detectors using the oscillation method. Separate low and high resolution passes were collected with different exposure times and detector distances and merged together in scaling, as the dynamic range of the diffraction data was larger than that of the detector. For the 274 K data, a Pilatus 6M pixel array detector (PAD) was used with shutterless data collection. Because of the very high dynamic range of the detector, a full dataset was collected in a single pass ∼5 minutes. The 274 K datasets were indexed and scaled using XDS^20^ while the 277K datasets were indexed and scaled and HKL2000^21^. The 100 K dataset was processed as previously described^13^.

### XFEL data collection and processing

For the XFEL experiment, microcrystals of ICH were grown by seeding. Initial seeds were obtained by pulverizing macroscopic crystals via vortexing with 0.5 mm stainless steel balls.for 5 minutes. A dilution of this seed stock was added to 31% PEG 3350, 250 mM MgCl_2_, 125 mM Tris-HCl pH=8.8, 2 mM DTT and then an equal volume of 40 mg/ml ICH was added and rapidly mixed. The mixture was incubated at room temperature with shaking for ∼20 minutes until microcrystal growth stopped. Samples were delivered to the beam using the concentric-flow microfluidic electrokinetic sample holder (coMESH) injector^22^ under atmospheric pressure conditions at room temperature and under normal atmosphere. The sample was loaded in a custom stainless steel sample reservoir. A Shimadzu LD20 HPLC pump hydraulically actuated a teflon plunger to advance the sample slurry. A stainless steel 20 &m frit filter (IDEX-HS) was placed after the reservoir in order to ensure smaller crystal sizes and mitigate clogging. The reservoir and filters were connected to the coMESH via a 100 &m x 160 &m x 1.5 m fused silica capillary (Molex). This capillary continued unobstructed through the center of the microfluidic tee (Labsmith) and terminated at 0 mm, 2 mm, or 200 mm recessed from the exit of a concentric 250 &m x 360 &m capillary, corresponding to APO, 15 s and 5 min time delays, respectively. The capillaries were optically aligned to obtain a 0 offset, then were recessed and measured externally to achieve the 2 mm and 200 mm recess. In the case of the longer 200 mm offset, an additional tee (IDEX-HS) was added to increase the length.

The outer line of the coMESH flowed the mother liquor, in the case of the APO structure, and flowed p-NPIC substrate for the 15 s and 5 min cases. The p-NPIC substrate was loaded into a 500 or 1000 &L gas tight syringe (Hamilton). The stainless steel, blunt tip removable needle was connected to a 250 &m x 360 &m x 1 m fused silica capillary (Molex) and connected to the side of the coMESH microfluidic tee junction for the APO and 15 s delay and into the side of the Idex tee for the 5 minute delay. The syringe was driven by a syringe pump (KDS Legato 200) at 1-2 &l/min while being charged at 3.1 kV (Stanford Research Systems, SRS PS300) at the wetted stainless steel needle.

The outer liquid focused the fluids ∼500 &m away from the outer capillary towards the XFEL focus, of approximately 3 &m at the MFX endstation. The charged meniscus was approximately 5 mm away from a grounded counter electrode to complete the electrokinetic focusing. The time delays were assumed to be sufficiently long as compared to the electrokinetic mixing phenomena and were determined simply by the time the bulk fluid would traverse the offset distance. The flow rates and voltages were held constant during each time point and the flow of the crystals were optically monitored with a 50x objective to assure minimal flow deviations. For the APO (no delay) time point, the flow was 0.5 &l/min for both the sample and outer flows. For the 15 s time delay, the capillaries were recessed 2 mm, and the flows were 1 and 2 &l/min for the sample and REACTANT flow rates, respectively. The combined bulk flows (1+2 = 3 &l/min) and the 2 mm offset dictated the delay time of 15 s. For the 5 min time delay, the capillaries were offset 200 mm and both flow rates were held at 1 &l/min, for a combined bulk flow of 2 &l/min. Overall, ∼0.5 mL of sample crystals and 0.5 mL of 3 mM p-NPIC were consumed to gather this data.

The XFEL diffraction data were processed with cctbx.xfel^23^, to identify 76,174 crystal hits for the APO data, 53,476 crystal hits for the 15 second delay data and 36,875 crystal hits for the 5 minute delay data. A small fraction of multiple lattice hits (<0.01% of total) were shown to have little impact on data quality when included in the final post-refinement steps. For the APO data, 23,669 frames were indexed and 21,834 of those had their intensities merged and integrated. For the 15s data, 26,507 frames were indexed and 24,286 of those had their intensities merged and integrated. For the 5min data, 16,893 frames were indexed and 14,483 of those had their intensities merged and integrated. To correct the intensity measurements and merge the data, postrefinement was carried out with three cycles of PRIME^24^ using model unit cell dimensions from a synchrotron wild type ICH structure^13^. The Lorentzian partiality model used parameters gamma_e = 0.001, frame_accept_min_cc = 0.60, uc_tolerance = 5, sigma_min = 2.5, and partiality_min = 0.2. The structure factor amplitudes and X-ray crystallographic data statistics for all datasets are provided in Table S1.

### Selection of mix-and-inject X-ray time delays

We collected absorbance spectra of ICH crystals in reservoir solution supplemented with 1mM p-napthyl isocyanide and 40% glycerol using UV-vis microspectrophotometry at SSRL BL 11-1 (Extended Data Figure S1). We freeze-trapped the reaction in the cryostream at fixed time-delays after adding substrate to the crystal before collecting a spectrum. The p-NPIC absorption maximum is ∼270 nm, while p-nitrophenyl formamide product absorption maximum is ∼315 nm on this instrument. The lack of an isosbestic point suggests an intermediate accumulated. At a time delay of 15s, the substrate peak at 270 nm had disappeared, while the product peak at 315 nm had not formed yet. At 5 minutes, substrate is exhausted while the product peak at 315 nm had fully formed, informing the timing for the XFEL experiment. The macroscopic (∼2×10^6^ µm^3^) crystals from which these spectra were collected were damaged by exposure to substrate and did not diffract X-rays well after substrate was introduced, precluding a cryotrapping X-ray diffraction experiment. In contrast, the microcrystals used for the XFEL experiment (∼2×10^3^ µm^3^) were not damaged by substrate, necessitating a serial X-ray crystallography approach.

### Crystallographic model refinement

All refinements were performed in PHENIX1.9 against structure factor intensities using individual anisotropic ADPs and riding hydrogen atoms^25^. Weights for the ADP refinement and relative weights for the X-ray and geometry term were optimized. After convergence of the isotropic ADP refinements, translation-libration-screw (TLS) refinements^26^ with automatic rigid body partitioning were performed in PHENIX. For the Cys101-SOH and G150A ICH synchrotron datasets, the alternate conformation for helix H (residues 148-173 in chain A) was modeled as a single occupancy group, reflecting the presumed correlated displacement of sidechain and backbone atoms. Other disordered regions were also built into multiple conformations manually in COOT^27^ if supported by 2mF_o_-DF_c_ and mF_o_-DF_c_ electron density maps. Occupancies of these groups were refined but constrained to sum to unity in PHENIX. Omit maps were calculated after removing the areas of interest from the model and refining for three macrocycles. Final models were validated using MolProbity^28^ and the validation tools in COOT and PHENIX. Model statistics are provided in Extended Data Table S2. Isomorphous difference maps (*F*_*o*_(15s) *– F*_*o*_(XFEL_APO_) and *F*_*o*_(SR) *– F*_*o*_(XFEL_APO_)) were calculated using data to the lowest high resolution of the data sets, 1.55 Å. Structure factor data and refined coordinates are available in the Protein Data Bank with the following accession codes: 6NI4, 6NI5, 6NI6, 6NI7, 6NI8, 6NI9, 6NJA, 6NPQ, XXXX, and XXXX.

### Synthesis of para-nitrophenyl isocyanide (p-NPIC)

ICH will accept diverse isocyanide substrates^12^. Due to its strong absorption in the visible range (%_max=_320 nm), para-nitrophenyl isocyanide (p-NPIC) was used here. The p-NPIC substrate is not commercially available and was synthesized by dehydration of the N-formamide precursor. To synthesize N-(4-Nitrophenyl)formamide, a mixture of amine (10 mmol) and formic acid (12 mmol) was stirred at 60 °C for 2h. After completion of the reaction (TLC), the mixture was diluted with CH_2_Cl_2_ (100 mL), washed with saturated solution of NaHCO_3_ and brine (100 mL), and then dried over anhydrous Na_2_SO_4_ and concentrated under reduced pressure. The residue was dissolved in hot acetone and crystallized, to yield N-(4-Nitrophenyl)formamide (1.58 g, 96%) as a yellow solid. The product was verified using ^1^H NMR, ^13^C NMR, IR, and Time of Flight (TOF) mass spectra. N-(4-Nitrophenyl)formamide (8.0 mmol) and triethylamine (56 mmol) were dissolved in CH_2_Cl_2_ (30 mL) and cooled to −10 °C. A solution of triphosgene (8.0 mmol) in CH_2_Cl_2_ (50 mL) was slowly added and stirring was continued at −10 °C for another 30 minutes. After completion of reaction (TLC), the reaction mixture was passed through a pad of silica gel and washed with CH_2_Cl_2_. The filtrate was dried over anhydrous Na_2_SO_4_, and then concentrated under reduced pressure. The residue was triturated with cold hexane to yield para-nitrophenyl isocyanide (1.36 g, 92%) as a yellow solid. ^1^H NMR, ^13^C NMR, IR, and Time of Flight (TOF) mass spectra were collected to verify the product structure.

### ICH enzyme kinetics

Steady state ICH rate measurements were initiated by the addition of ICH (final concentration of 1 &M) to freshly prepared p-NPIC solutions ranging from 0-60 &m in reaction buffer (100 mM KPO_4_ pH=7.0, 50 mM KCl, and 20% DMSO). p-NPIC was diluted from a 0.5 M stock solution in dimethyl sulfoxide (DMSO), which was stored at −80 °C and protected from light. A 200 &L reaction was maintained at 25 °C in a Peltier-thermostatted cuvette holder. The formation of the product, para-nitrophenyl formamide (p-NPF) was monitored at its absorption maximum of 320 nm for two minutes using a UV-Vis Cary 50 Spectrophotometer (Varian, Palo Alto, CA). A linear increase in A_320_ was verified and used to calculate initial velocities. Isocyanides can spontaneously hydrolyze slowly in aqueous solutions, with the rate increasing as pH is lowered. To ensure that all measured product formation was due to ICH catalysis, the rate of spontaneous p-NPIC hydrolysis was measured without added ICH. These values were near the noise level of the spectrophotometer and were subtracted from the raw rate measurements before fitting the Michaelis-Menten model. The extinction coefficient at 320 nm for p-NPF was determined by using ICH to convert known concentrations of p-NPIC to p-NPF, followed by measuring the absorbance at 320 nm. The slope of the resulting standard curve was defined as the extinction coefficient of p-NPF at 320 nm; ‘_320_=1.33×10^4^ M^-1^ cm^-1^. This p-NPF ‘_320_ value was used to convert the measured rates from A_320_/sec to [p-NPF]/sec. All data were measured in triplicate or greater and mean values and standard deviations were plotted and fitted using the Michaelis-Menten model as implemented in Prism (GraphPad Software, San Diego, CA). Reported K_m_ and k_cat_ values and their associated errors are given in Extended Data Figure S8.

Pre-steady state ICH kinetics were measured at 25 °C using a Hi-Tech KinetAsyst stopped flow device (TgK Scientific, Bradford-on-Avon, United Kingdom). Data were collected for each sample for two seconds (instrument deadtime is 20 ms) in triplicate. The final enzyme concentration after mixing was 10 μM and final p-NPIC concentrations were 20, 40, 80, 120, 160, and 320 μM. Product evolution was monitored at 320 nm using a photodiode array detector. Kinetic Studio software (TgK Scientific, United Kingdom) was used to analyze the kinetic data and to fit a mixed model containing a single exponential burst with a linear steady state component: -Aexp(-kt) + mt + C. In this equation, t is time, k is the burst phase rate constant, m is the linear phase (steady-state) rate, A is the amplitude of the burst component, and C is a baseline offset constant. The linear slope m was used to calculate steady state turnover numbers, which agree well with k_obs_ values obtained from steady state kinetic measurements at comparable substrate concentrations. Single turnover experiments were also performed using final concentrations of 40 μM enzyme and 20 μM p-NPIC after manual mixing. Spectra were collected using a Cary 50 spectrophotometer (Varian, Palo Alto, CA, USA). Spectra were collected every 10 seconds with a deadtime of ∼30 seconds after manual addition of enzyme.

### Molecular Dynamics Simulations

An ICH dimer was protonated at pH 7.0 and minimized under the OPLS3 force field using the Schrödinger 2018-3 software suite [Schrödinger Suite 2018-3 Protein Preparation Wizard; Schrödinger, LLC, New York, NY, 2016]. ICH crystals were parameterized using Amber^29^. A simulation cell with unit cell dimensions and P2_1_ symmetry of the crystal structure was created and replicated to obtain a 2×2×2 supercell containing 16 ICH dimers to minimize boundary artefacts in the simulations. 200 mM MgCl_2_ and 100mM Tris-HCl were added to the simulation cell. While the crystallization conditions also included PEG3350 and DTT, those were omitted from the simulation owing to the large size of PEG and the low concentration (2mM) of DTT. The system was electrostatically neutralized by adding Na^+^ ions. SPC/E waters were added to obtain a pressure of approximately 1bar. Simulations of ‘XFEL conditions’ were started from the XFEL crystal structure, with C101 in the thiolate state. By contrast, the synchrotron structure was simulated by modifying C101 to Cys-sulfenate (Cys-SOH) in the XFEL structure and removing the catalytic water near Asp17. Partial charges for Cys-SOH were determined with HF/6-31G* basis set and the AM1-BCC method in Antechamber^30^. The systems were minimized with steepest descent and conjugate gradient algorithms by gradually reducing constraints on the protein atoms, and heated (NVT) to 300K over 10ps. The time step was set to 1 fs for heating and the initial phase of equilibration. Production runs of 1 μs ICH crystal NVT ensembles were carried out at 300K with openMM^31^ on NVIDIA K80 and P100 GPUs, with a Langevin thermostat (friction coefficient 1.0), an integration timestep of 2 fs, and non-bonded cutoffs of 1 nm. In total, we obtained 4 μs of simulation time for ICH.

The distance of the I152_H_-C101_SG_ hydrogen bond was calculated for each protomer at time intervals of 500 ps over the course of the simulations. The average I152_H_-C101_SG_ distance was obtained by averaging over the protomers (bold line, Figure 3B), and then averaging over time. The shifts of helix H were calculated at time intervals of 500 ps by first aligning each protomer to the first frame using all backbone (heavy) atoms giving an RMSD_REF_, and then calculating the backbone (heavy) atom RMSD to the first frame over residues 152 to 166 giving RMSD_HELIX_. We then reported the fraction RMSD_HELIX_/ RMSD_REF_. While RSMD strongly depends on the length of the fragment and flexible loops, generally RMSD_HELIX_/ RMSD_REF_ > 1 indicates that the helix shifts more than the rest of the protomer. The average shift was obtained by averaging over the protomers (bold line, Figure 3C), and then averaging over time.

## References

1 Danyal, K., Mayweather, D., Dean, D. R., Seefeldt, L. C. & Hoffman, B. M. Conformational gating of electron transfer from the nitrogenase Fe protein to MoFe protein. J. Am. Chem. Soc. 132, 6894–6895, (2010).

2 Henzler-Wildman, K. A. et al. Intrinsic motions along an enzymatic reaction trajectory. Nature 450, 838–844, (2007).

3 Aviram, H. Y. et al. Direct observation of ultrafast large-scale dynamics of an enzyme under turnover conditions. Proc. Natl. Acad. Sci. USA 115, 3243–3248, (2018).

4 Kim, T. H. et al. The role of dimer asymmetry and protomer dynamics in enzyme catalysis. Science 355, (2017).

5 Hammes, G. G. Multiple conformational changes in enzyme catalysis. Biochemistry 41, 8221–8228, (2002).

6 Kale, S. et al. Efficient coupling of catalysis and dynamics in the E1 component of Escherichia coli pyruvate dehydrogenase multienzyme complex. Proc. Natl. Acad. Sci. USA 105, 1158–1163, (2008).

7 Kupitz, C. et al. Structural enzymology using X-ray free electron lasers. Struct. Dyn. 4, 044003, (2017).

8 Stagno, J. R. et al. Structures of riboswitch RNA reaction states by mix-and-inject XFEL serial crystallography. Nature 541, 242–246, doi:10.1038/nature20599 (2017).

9 Tosha, T. et al. Capturing an initial intermediate during the P450nor enzymatic reaction using time-resolved XFEL crystallography and caged-substrate. Nat. Commun. 8, 1585, (2017).

10 Olmos, J. L., Jr. et al. Enzyme intermediates captured “on the fly” by mix-and-inject serial crystallography. BMC Biol. 16, 59, (2018).

11 Spence, J. C. H. XFELs for structure and dynamics in biology. IUCr. 4, 322–339, (2017).

12 Goda, M., Hashimoto, Y., Shimizu, S. & Kobayashi, M. Discovery of a novel enzyme, isonitrile hydratase, involved in nitrogen-carbon triple bond cleavage. J. Biol. Chem. 276, 23480–23485, (2001).

13 Lakshminarasimhan, M., Madzelan, P., Nan, R., Milkovic, N. M. & Wilson, M. A. Evolution of new enzymatic function by structural modulation of cysteine reactivity in Pseudomonas fluorescens isocyanide hydratase. J. Biol. Chem. 285, 29651–29661, (2010).

14 Brereton, A. E. & Karplus, P. A. Native proteins trap high-energy transit conformations. Sci. Adv. 1, e1501188, (2015).

15 Chapman, H. N., Caleman, C. & Timneanu, N. Diffraction before destruction. Philos Trans. R. Soc. Lond. B Biol. Sci. 369, 20130313, (2014).

16 van den Bedem, H., Dhanik, A., Latombe, J. C. & Deacon, A. M. Modeling discrete heterogeneity in X-ray diffraction data by fitting multi-conformers. Acta crystallogr. D, Biol. Crystallogr. 65, 1107–1117, (2009).

17 van den Bedem, H., Bhabha, G., Yang, K., Wright, P. E. & Fraser, J. S. Automated identification of functional dynamic contact networks from X-ray crystallography. Nature methods 10, 896–902, (2013).

18 Budday, D., Leyendecker, S. & van den Bedem, H. Kinematic Flexibility Analysis: Hydrogen Bonding Patterns Impart a Spatial Hierarchy of Protein Motion. J. Chem. Inf. Model 58, 2108–2122, (2018).

19 van den Bedem, H. & Fraser, J. S. Integrative, dynamic structural biology at atomic resolution--it’s about time. Nat. methods 12, 307–318, doi:10.1038/nmeth.3324 (2015).

20 Kabsch, W. Integration, scaling, space-group assignment and post-refinement. Acta crystallogr. D, Biol. crystallogr. 66, 133–144, (2010).

21 Otwinowski, Z. & Minor, W. Processing of X-ray diffraction data collected in oscillation mode. Macromolecular Crystallography, Pt A 276, 307–326 (1997).

22 Sierra, R. G. et al. Concentric-flow electrokinetic injector enables serial crystallography of ribosome and photosystem II. Nat. methods 13, 59–62, (2016).

23 Sauter, N. K., Hattne, J., Grosse-Kunstleve, R. W. & Echols, N. New Python-based methods for data processing. Acta crystallogr. D, Biol.l crystallogr. 69, 1274–1282, (2013).

24 Uervirojnangkoorn, M. et al. Enabling X-ray free electron laser crystallography for challenging biological systems from a limited number of crystals. eLife 4, (2015).

25 Adams, P. D. et al. PHENIX: a comprehensive Python-based system for macromolecular structure solution. Acta crystallogr. D, Biol. crystallogr.y 66, 213–221, (2010).

26 Schomaker, V. & Trueblood, K. N. On Rigid-Body Motion of Molecules in Crystals. Acta Crystall B-Stru B 24, 63-+, (1968).

27 Emsley, P. & Cowtan, K. Coot: model-building tools for molecular graphics. Acta crystallogr. D, Biol. crystallogr. 60, 2126–2132, (2004).

28 Davis, I. W. et al. MolProbity: all-atom contacts and structure validation for proteins and nucleic acids. Nucleic Acids Res. 35, W375–383, (2007).

29 Case, D. A. et al. The Amber biomolecular simulation programs. J. Comput. Chem. 26, 1668–1688, (2005).

30 Wang, J., Wang, W., Kollman, P. A. & Case, D. A. Automatic atom type and bond type perception in molecular mechanical calculations. J. Mol. Graph. Model 25, 247–260, (2006).

31 Eastman, P. & Pande, V. S. OpenMM: A Hardware Independent Framework for Molecular Simulations. Comput. Sci. Eng. 12, 34–39, (2015).

