## Extended Data for "Mix-and-inject XFEL crystallography reveals gated conformational dynamics during enzyme catalysis"

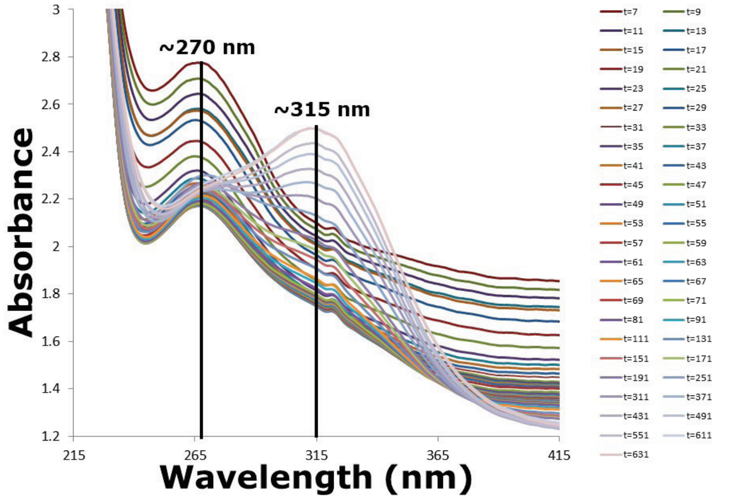


**Figure S1.** ***In crystallo*** **UV-visible spectrophotometry of ICH catalysis.** A single crystal of ICH was mixed with 1 mM p-NPIC, incubated for the indicated times in seconds, mounted and cryocooled to 100 K in the cryostream. Absorbance spectra were collected using the inline spectrophotometer at BL 11-1 at the Stanford Synchrotron Radiation Lightsource.

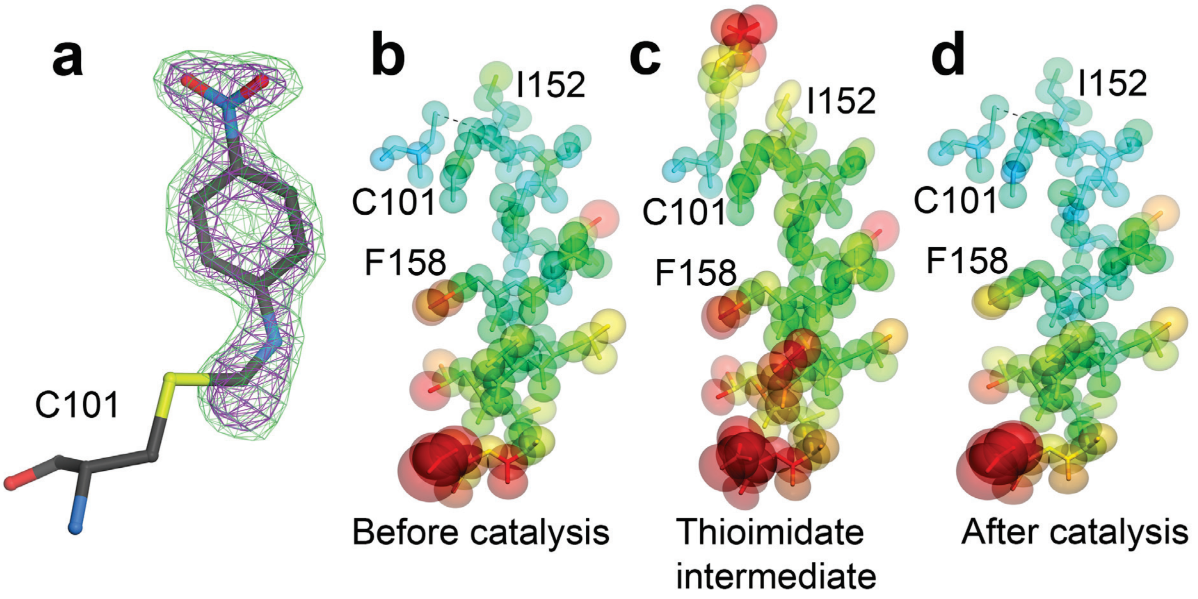


**Figure S2. Observation of thioimidate intermediate formation and enhancement of ICH helix mobility.** In (a), omit $mF_o$-$DF_c$ electron density is shown at 3.0 rmsd (green) and 5.0 rmsd (purple), unambiguously demonstrating presnece of the catalytic intermediate. In (b-d), anisotropic atomic displacement parameters (ADPs) from TLS models refined against the XFEL datasets are shown at the 75% probability level and colored according to $B_{eq}$ value, from blue (7 Å$^2$) to red (30 Å$^2$). Starting with the resting enzyme before substrate is introduced (b), helical mobility is transiently elevated upon thioimidate intermediate formation during catalysis (c) and then is reduced again upon completion of catalysis (d) once substrate is consumed 5 minutes after mixing.

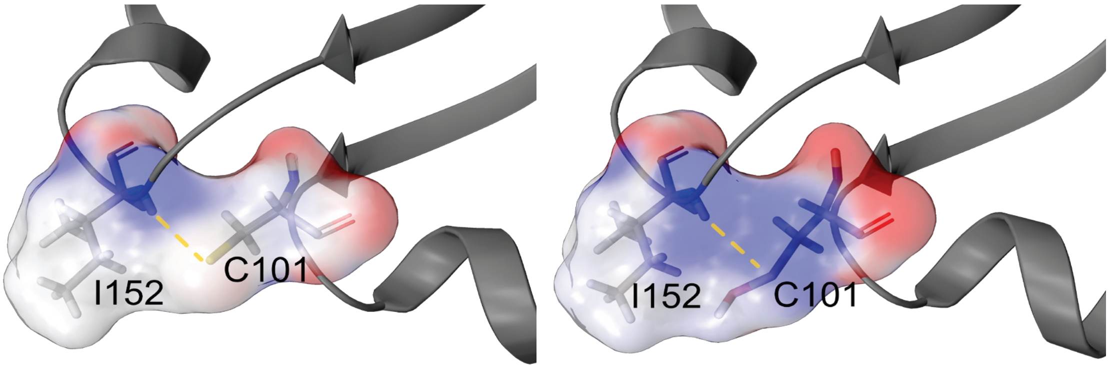


**Figure S3. Cys101 oxidation leads to a weakening of the Ile152-Cys101 H-bond.** Electrostatic Poisson-Boltzmann surfaces (red negative, blue positive charge) calculated from the Cys101-Ile152 (A) and Cys-SOH-Ile152 (B) environments. Cysteine photooxidation neutralizes the negative charge of the sulfur atom, weakening the N-H … S hydrogen bond. We calculated a reduction in the Cys101-Ile152 hydrogen bond energy from -2.2 kcal/mol under reduced conditions (i.e. with a thiolate acceptor) to -0.91 kcal/mol upon Cys101-SOH formation [Methods (3)]

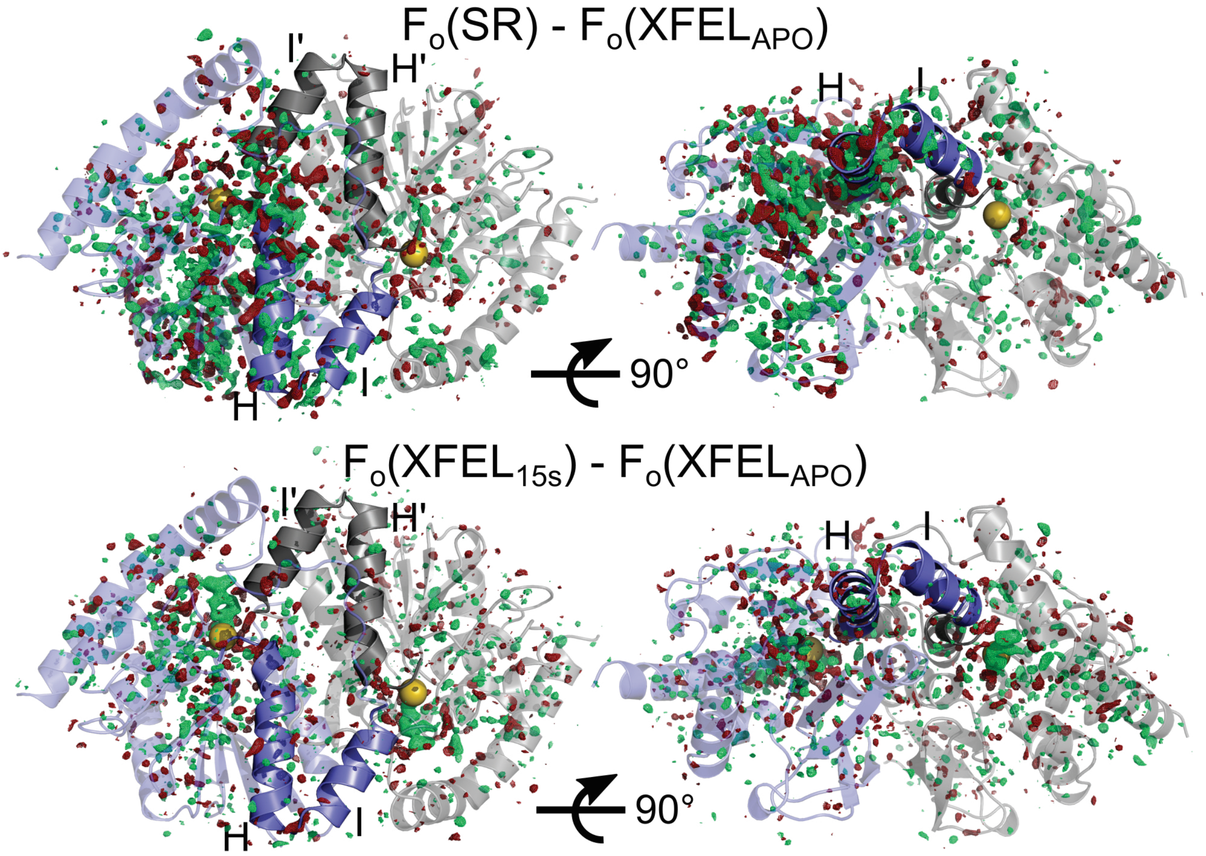


**Figure S4. Isomorphous $F_o$(SR) – $F_o$($XFEL_{APO}$) (top panels) and $F_o$($XFEL_{15s}$) – $F_o$($XFEL_{APO}$) (bottom panels) difference maps at 2.75 rmsd.** Difference features (green, positive; red, negative) are non-uniformly distributed in both maps, radiating out from the site of the catalytic nucleophile (yellow spheres). This is more pronounced in the A protomer (slate) than the B protomer (grey) in the $F_o$(SR) – $F_o$($XFEL_{APO}$) map (top panels). Difference features generally show the same, but weaker pattern in the $F_o$($XFEL_{15s}$) – $F_o$($XFEL_{APO}$) map. The maps are phased with the APO XFEL structure.

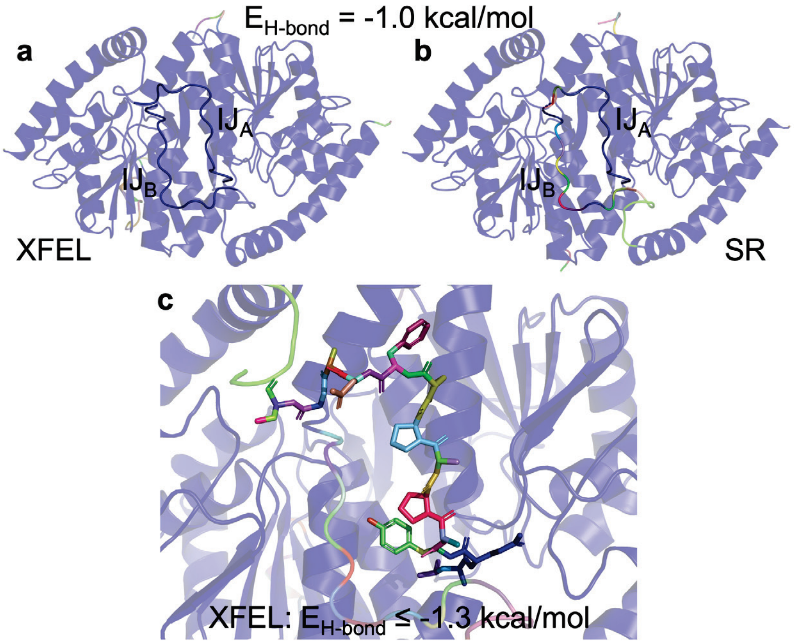


**Figure S5. Rigidity Analysis.** Individual rigid clusters are color-coded. In both the XFEL (panel **a**) and SR (panel **b**) structures, the backbone is largely rigidified (blue color). **a**) ICH XFEL structure. The $IJ_A$ and $IJ_B$ linkers are rigid in the XFEL structure when H-bonds stronger than -1 kcal/mol are included in the analysis, with the C101:S-I152:NH bond constraint in both protomers. **b**) ICH SR structure. The $IJ_B$ linker, which contacts helix H, becomes flexible in the SR structure with the shifted conformation of helix H. **c**) When H-bonds weaker than -1.3 kcal/mol are excluded, both linkers $IJ_A$ and $IJ_B$ become flexible. Interestingly, these are the first structural elements of ICH to melt when weaker H-bonds are omitted.

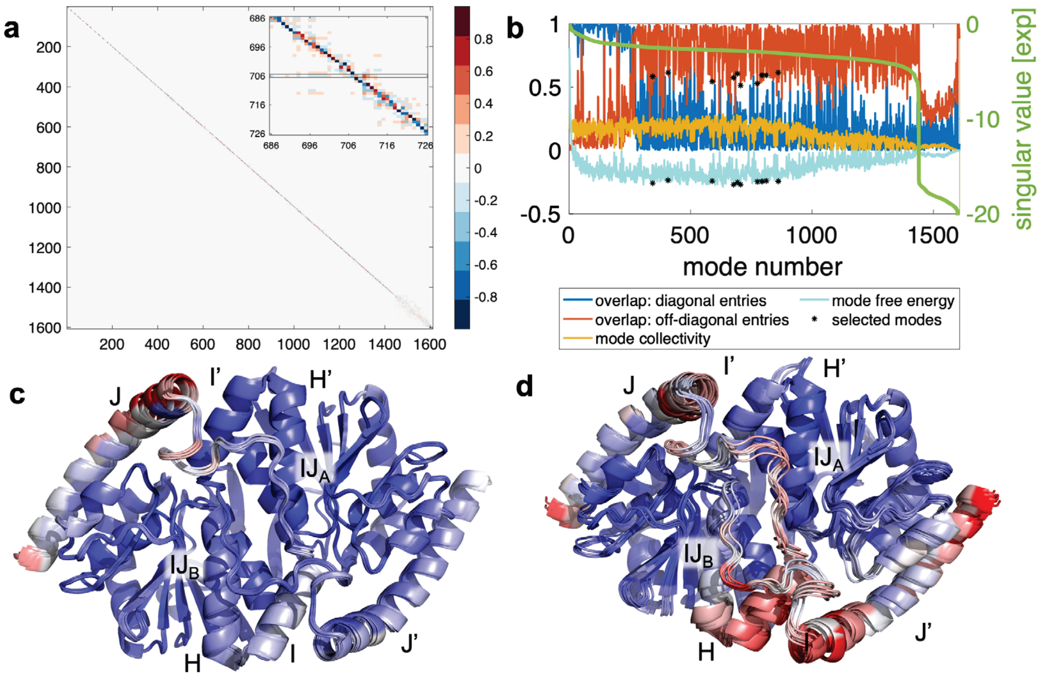


**Figure S6**. **a**) Visualization of the matrix product $\boldsymbol{D} = \boldsymbol{V}_B^T \boldsymbol{V}_{A\&B}$. The matrix $\boldsymbol{D}$ reports on the mode overlap when the Cys101 hydrogen bond is present only in the B protomer (vertical) vs. in both protomers (horizontal) of the XFEL structure. Motion vector dot products are color coded between -1 and 1 (color bar). The inset highlights an area around one of the modes with lowest mode-specific free energy and least overlap to any other mode (mode 706, insert). **b**) Motion modes that are least similar between the XFEL and SR structures. The mode-specific free energy (teal) is the difference between internal energy perturbation (singular values in green line) and conformational entropy (mode collectivity, yellow) computed as the normalized exponential of the Shannon entropy. From a set of 100 modes with lowest free energy (teal), we select the top ten modes with least overlap (red and blue) to any other mode. For each of these modes, the two stars indicate their mode-specific free energy and their maximum diagonal (blue) or off-diagonal (red) overlap to any other mode. **c-d**) Variations in ICH motional spectrum and inter-protomeric exchange upon Cys101 hydrogen-bond modification. **c**) Conformational ensemble of motion modes in Kinematic Flexibility Analysis enriched after the C101$_{SG}$-I152$_H$ H-bond in the A protomer is disrupted. Consistent with the MD simulations, the IJ linkers in both conformers, which are in contact with helices H, show increased conformational dynamics, especially near both active sites. Motion modes associated with helix H and, in particular, I, and helices J and J' are also enriched. **d**) For comparison, we also selected ten randomly selected KFA orthogonal motion modes with mode numbers < 300 or > 1,000 (elevated mode-specific free energies) and computed their associated motions with the same step size. These motions are less focused, and engage the entire protein. The conformational ensembles are projected onto the XFEL structure and colored by RMSF, increasing from blue to red.

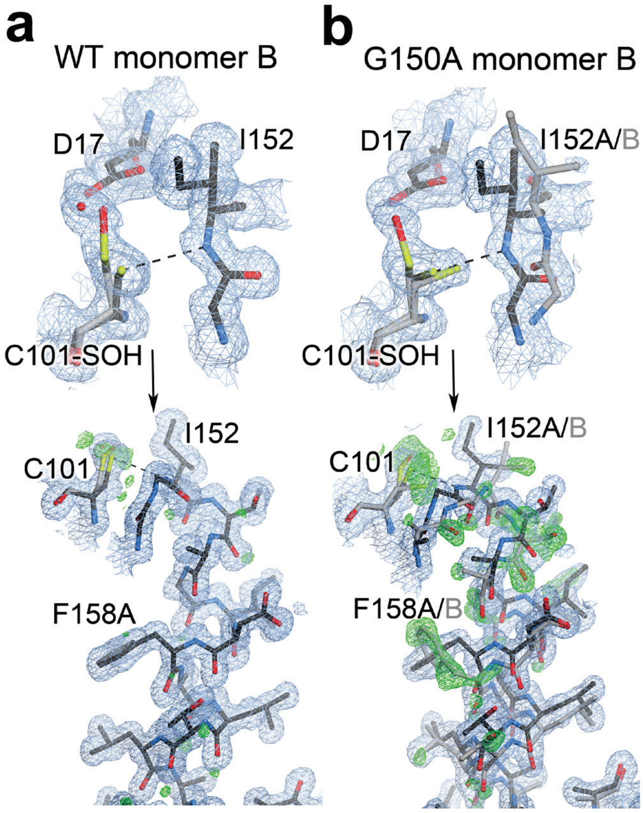


**Figure S7: Helical mobility is asymmetric in wild-type but not G150A ICH.** The top panels show the environment of Cys101 in the second protomer ("monomer B") of wild-type (WT) and G150A ICH. $2mF_o$-$DF_c$ electron density is contoured at 0.7 RMSD (blue) and the hydrogen bond between the peptide backbone of Ile152 and Cys101 is shown in a dotted line. The lower panels show the helix in its strained (black) and relaxed, shifted conformations (grey). $2mF_o$-$DF_c$ electron density is contoured at 0.8 RMSD (blue) and omit $mF_o$-$DF_c$ electron density for the shifted helical conformation is contoured at 3.0 rmsd (green). Wild-type ICH does not show strong evidence of a second conformation, even though Cys101 has been partially photooxidized to Cys101-SOH (top). In contrast, the helix in monomer B of G150A ICH shows strong evidence of a second, relaxed conformation, similar to monomer A.

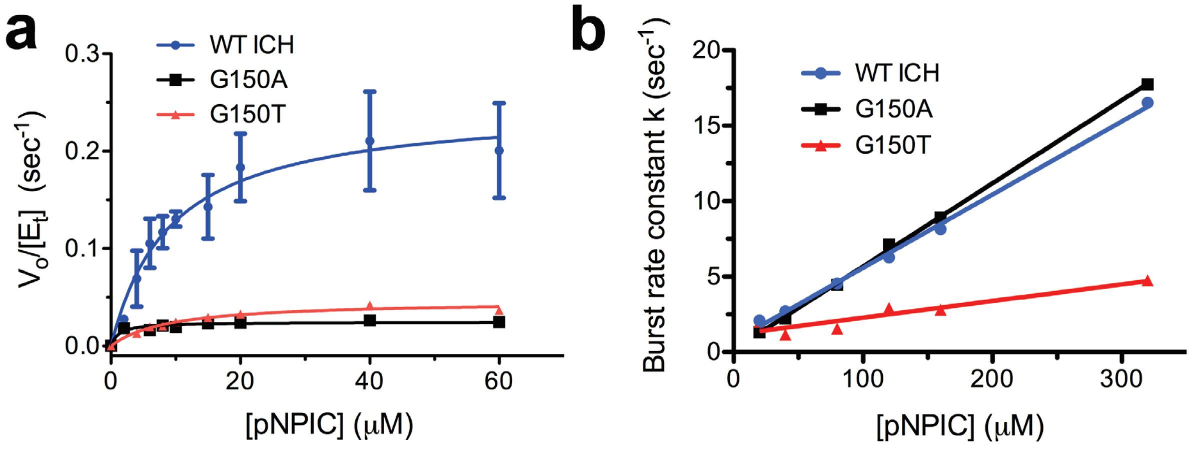


**Figure S8:** Steady-state (a) and pre-steady state (b) enzyme kinetics of wild-type (WT; blue circles), G105A (black squares), and G150T (red triangles) ICH. Both the G150A and G150T mutations result in similar decreases in steady-state kinetics compared to WT enzyme ($k_{cat}$(WT)=0.248±0.031 $s^{-1}$, $k_{cat}/K_M$(WT)= 2.68x10$^4$ $M^{-1}s^{-1}$; $k_{cat}$(G150A)=0.025±0.002 $s^{-1}$, $k_{cat}/K_M$(G150A)=2.07x10$^4$ $M^{-1}s^{-1}$; $k_{cat}$(G150T)=0.046±0.003 $s^{-1}$, $k_{cat}/K_M$(G150T)= 5.21x10$^3$ $M^{-1}s^{-1}$). (b) The burst rate constant in pre-steady state kinetics is linearly dependent on substrate concentration. This linear dependence indicates a second order rate process during the burst phase, consistent with thioimidate intermediate formation. WT and G150A have similar second order burst rate constants (k(WT)=4.85x10$^4$±943 $M^{-1}s^{-1}$; k(G150A)= 5.51x10$^4$±557 $M^{-1}s^{-1}$), but G150T ICH is lower (k(G150T)= 1.11x10$^4$±999 $M^{-1}s^{-1}$).

**Table S1: Crystallographic Data Statistics**

| Sample | WT ICH RT[1] (less oxidized) | WT ICH RT[1] (more oxidized) | G150A ICH | G150T ICH | WT ICH XFEL: Apo[1] | WT ICH XFEL: 15 sec Thioimidate[1] | WT ICH XFEL: 5 min[1] | WT ICH cryo |
|---|---|---|---|---|---|---|---|---|
| Diffraction source | SSRL 12-2 | SSRL 7-1 | SSRL 12-2 | SSRL 7-1 | LCLS MFX | LCLS MFX | LCLS MFX | APS 14BM-C |
| Wavelength (Å) | 0.827 | 0.975 | 0.827 | 0.975 | 1.305 | 1.305 | 1.305 | 0.900 |
| Temperature (K) | 274 | 277 | 274 | 277 | 298 | 298 | 298 | 100 |
| Detector | Pilatus 6M | ADSC Q315 | Pilatus 6M | ADSC Q315 | Rayonix MX170-HS | Rayonix MX170-HS | Rayonix MX170-HS | ADSC Q315 |
| Space group | $P2_1$ | $P2_1$ | $P2_1$ | C2 | $P2_1$ | $P2_1$ | $P2_1$ | $P2_1$ |
| a, b, c (Å) | 57.24 58.03 69.09 | 57.15 57.97 69.09 | 57.04 57.90 69.11 | 72.11 59.74 56.13 | 56.88 57.68 68.78 | 56.77 57.42 68.79 | 56.81 57.64 68.76 | 56.58 56.47 68.23 |
| α, β, γ (°) | 90.00 112.83 90.00 | 90.00 112.77 90.00 | 90.00 112.55 90.00 | 90.00 115.86 90.00 | 90.00 112.74 90.00 | 90.00 112.74 90.00 | 90.00 112.74 90.00 | 90.00 112.49 90.00 |
| Mosaicity (°) | 0.08 | 0.08 | 0.08 | 0.07 | N/A[5] | N/A[5] | N/A[5] | 0.27 |
| Resolution range (Å) | 39.04-1.20 (1.22-1.20) | 42.87-1.15 (1.17-1.15) | 38.96-1.30 (1.32-1.30) | 35.45-1.10 (1.12-1.10) | 20.13-1.55 (1.57-1.55) | 20.09-1.55 (1.57-1.55) | 20.1-1.55 (1.57-1.55) | 37-1.05 (1.09-1.05) |
| Total no. of observations | 363112 (16358) | 517060 (15585) | 327591 (13542) | 224032 (2017) | 3032726 (18818) | 3185312 (16342) | 2648001 (38049) | 1077211 (79769) |
| No. of unique observations | 126373 (5922) | 145885 (6657) | 100249 (4568) | 84423 (1216) | 60075 (2983) | 60053 (2970) | 60082 (2988) | 179423 (17341) |
| Completeness (%) | 97.4 (92.7) | 98.9 (91.2) | 98.1 (90.1) | 97.2 (95.9) | 99.9 (99.8) | 99.9 (99.4) | 99.9 (100) | 97.1 (94.2) |
| Multiplicity | 2.9 (2.8) | 3.5 (2.3) | 3.3 (3.0) | 2.7 (2.0) | 50.5 (6.3) | 53.0 (5.5) | 44.1 (12.7) | 6.0 (4.6) |
| ⟨I/σ(I)⟩ | 10.4(1.8) | 12.5 (0.8) | 9.1 (1.2) | 9.7 (1.4) | 60.5 (2.0) | 58.4 (1.8) | 73.1 (3.4) | 22.8 (2.1) |
| $CC_{1/2}$[2] | 0.991 (0.225) | 0.999 (0.488) | 0.996 (0.318) | 0.999 (0.690) | 0.962 (0.16) | 0.963 (0.14) | 0.937 (0.36) | N/A[4] |
| $R_{meas}$ ($R_{split}$ for XFEL data)[3] | 0.072 (1.689) | 0.059 (0.959) | 0.066 (1.513) | 0.045 (0.545) | 0.196 (0.862) | 0.197 (0.905) | 0.232 (0.658) | 0.07 (0.70) |

Values for the highest resoltions bin shown in parenthesis
[1] RT, Room temperature synchrotron radiation collection; XFEL, X-ray free electron laser
[2] $CC_{1/2}$ [1] was used to determine the high resolution cutoff.
[3] $R_{split}$ is provided only for XFEL serial crystallographic data
[4] Dataset from PDB 3NON, processed before the introduction of $CC_{1/2}$
[5] N/A=not applicable for serial crystallographic data

**Table S2: Crystallographic Refinement Statistics**

| Model | WT ICH RT[1] (less oxidized) | WT ICH RT[1] (more oxidized) | G150A ICH | G150T ICH | WT ICH XFEL: Apo[1] | WT ICH XFEL: 15 sec Thioimidate[1] | WT ICH XFEL: 5 min[1] | WT ICH RT[1] (less oxidized) qFit model | WT ICH cryo with helix disorder |
|---|---|---|---|---|---|---|---|---|---|
| PDB code | 6NI6 | 6NI7 | 6NI5 | 6NI4 | 6NPQ | XXXX | XXXX | 6NI9 | 6NJA |
| Temperature (K) | 274 | 277 | 274 | 277 | 298 | 298 | 298 | 274 | 100 |
| Refinement program | PHENIX 1.9 | PHENIX 1.9 | PHENIX 1.9 | PHENIX 1.9 | PHENIX 1.9 | PHENIX 1.9 | PHENIX 1.9 | PHENIX 1.9 | PHENIX 1.9 |
| Resolution range (Å) | 31.84-1.20 | 38.53-1.15 | 33.56-1.30 | 35.14-1.10 | 20.13-1.55 | 20.09-1.55 | 17.46-1.55 | 31.84-1.20 | 37.82-1.05 |
| Completeness (%) | 95.17 | 98.87 | 97.83 | 97.16 | 99.93 | 99.86 | 99.96 | 95.00 | 97.11 |
| No. of reflections | 123572 | 145829 | 99966 | 84275 | 59628 | 59214 | 59498 | 123572 | 179399 |
| No. of reflections, test set | 6203 | 7300 | 4970 | 4186 | 2002 | 1986 | 1995 | 6203 | 8979 |
| $R_{work}$ | 0.1140 (0.2281) | 0.1128 (0.2633) | 0.1192 (0.2745) | 0.1098 (0.2294) | 0.1560 (0.3075) | 0.1636 (0.3244) | 0.1727 (0.2777) | 0.1078 (0.2207) | 0.1170 (0.2105) |
| $R_{free}$ | 0.1398 (0.2611) | 0.1353 (0.2607) | 0.1519 (0.3159) | 0.1283 (0.2332) | 0.1864 (0.3244) | 0.1906 (0.3177) | 0.1918 (0.2836) | 0.1402 (0.2628) | 0.1342 (0.2487) |
| No. of non-H atoms | | | | | | | | | |
| Protein | 4486 | 4446 | 4892 | 1868 | 3475 | 3449 | 3500 | 6192 | 4551 |
| Water | 342 | 323 | 323 | 149 | 298 | 271 | 272 | 382 | 484 |
| Total | 4828 | 4769 | 5215 | 2017 | 3773 | 3720 | 3772 | 6574 | 5035 |
| Average R.M.S. deviations | | | | | | | | | |
| Bonds (Å) | 0.007 | 0.008 | 0.014 | 0.011 | 0.007 | 0.004 | 0.005 | 0.011 | 0.009 |
| Angles (°) | 0.909 | 1.020 | 1.290 | 1.376 | 0.850 | 0.729 | 0.801 | 1.399 | 1.111 |
| Average B factors ($<B_{iso}>$, Å$^2$) | | | | | | | | | |
| Protein | 17.34 | 19.57 | 21.12 | 17.21 | 24.18 | 24.97 | 25.78 | 15.92 | 11.91 |
| Water | 36.77 | 37.40 | 39.73 | 31.30 | 38.83 | 38.61 | 44.80 | 36.74 | 26.43 |
| Average ADP anisotropy[2] | | | | | | | | | |
| Protein | 0.396 | 0.481 | 0.400 | 0.502 | 0.599 | 0.553 | 0.626 | 0.394 | 0.407 |
| Water | 0.327 | 0.415 | 0.353 | 0.439 | 1 | 1 | 1 | 0.354 | 0.408 |
| MolProbity clashscore | 1.1 | 2.2 | 4.1 | 0.8 | 2.0 | 1.6 | 1.3 | 2.4 | 3.2 |
| Ramachandran plot | | | | | | | | | |
| Outliers (%) | 0.5 | 0.3 | 0.5 | 0.0 | 0.4 | 0.4 | 0.4 | 0.3 | 0.3 |
| Allowed (%) | 1.4 | 1.4 | 2.0 | 1.2 | 1.3 | 1.3 | 1.8 | 1.1 | 1.9 |
| Favored (%) | 98.1 | 98.3 | 97.5 | 98.8 | 98.2 | 98.2 | 97.8 | 98.6 | 97.8 |

Values for the highest resoltions bin shown in parenthesis

[1] RT, Room temperature synchrotron radiation collection; XFEL, X-ray free electron laser

[2] Anisotropy is defined as the ratio of the smallest to largest eigenvalue of the ADP tensor

### Extended Results

**Propagation of cysteine-gated conformational changes across the ICH dimer.** To characterize the extent of the conformational response of the entire ICH dimer to the modification of Cys101, we calculated an $F_o$(SR) – $F_o$($XFEL_{APO}$) isomorphous difference map, phased with a structural model obtained from the XFEL data set. The $F_o$(SR) – $F_o$($XFEL_{APO}$) isomorphous difference map provides an unbiased view of differences in molecular conformation between two data sets, reporting specifically on changes in ICH that occur in response to oxidation of Cys101. A $F_o$(15s) – $F_o$($XFEL_{APO}$) map calculated using the thioimidate dataset after 15 sec of mixing and the dataset prior to introduction of subtrate (Apo) reveals similar features to those seen in the $F_o$(SR) – $F_o$($XFEL_{APO}$) map (Extended Data Figure S4), signifying that the widespread dynamic response observed in $F_o$(SR) – $F_o$($XFEL_{APO}$) difference map is due to Cys101 oxidation.

We further examined the dynamical communication across the dimer interface in ICH using CONTACT network analysis. CONTACT elucidates pathways of collective amino acid main- and sidechain displacements through mapping van der Waals conflicts that would result from sidechain conformational disorder if correlated motions are not considered [2]. CONTACT identified a large network of correlated residues in protomer A (with the mobile helix) that connects with a smaller network in protomer B (Figure 4D), corroborating the isomorphous difference map. The key residues in CONTACT analysis that bridge the dimer interface are Tyr181 and Thr153 (Figure 4E), which are also key residues identified in the isomorphous difference map.

We computed rigid cluster decompositions of ICH corresponding to the unshifted (XFEL) and shifted (SR) conformations. We removed all alternate conformations in the three crystal structures, retaining only conformations corresponding to the unshifted state in the XFEL structure, and the shifted state in the SR structure. Waters were retained. These structures were protonated at pH 7.0 and minimized under the OPLS3 force field using the Schrödinger 2018-3 software suite [Schrödinger Suite 2018-3 Protein Preparation Wizard; Schrödinger, LLC, New York, NY, 2016]. We then used our KGS software suite[3] to carry out rigidity analysis, using hydrogen bonds stronger than -1 kcal/mol and hydrophobic interactions as constraints. In the XFEL structure, the Cys101-Ile152 H-bond was added as a constraint in both protomers. In the SR structure, the Cys101-Ile152 H-bond was included only in the B protomer.

All structures show a largely rigid core (Extended Data Figure S5), although the largest cluster (blue) is slightly bigger in the SR structure than in the XFEL structure. The IJ linkers (opaque) are rigid in the XFEL structure (Extended Data Figure S5A), while linker $IJ_B$ melted in the SR structure, concomitant with the shifted helix $H_A$ (Extended Data Figure S5B), signifying that the loss of a non-covalent interaction in response to Cys101 oxidation allosterically propagates across the protein. Notably, we found that the IJ linkers are the first major structural elements to melt once weaker H-bonds are excluded from the analysis (Extended Data Figure S5C).

To understand how the active site Cys101-Ile152 hydrogen bond modulates conformational dynamics, we used Kinematic Flexibility Analysis (KFA[4]). KFA represents a molecule as a kinematic linkage with dihedral degrees of freedom and hydrogen bonds and hydrophobic contacts as constraints. Conventional rigidity analysis[3] of ICH with KFA revealed that changes in protein flexibility between the unshifted and shifted conformations are concentrated on the IJ linkers (Figure S5). In contrast to traditional molecular rigidity analysis, KFA can provide an explicit basis for orthogonal protein motion modes coupled to energetic penalties incurred by perturbing the constraint network. KFA can rank-order protein motions by the magnitude of free energy changes calculated from non-covalent interactions and molecular rigidity (see details below). We analyzed how motion modes corresponding to the lowest free energies in the structure without Cys101 modification change when the H-bond is intact in both A and B protomers (C101-I152$_{A\&B}$) to when this H-bond is disrupted in protomer A (C101-I152$_{B}$). These altered motion modes are the ones most affected by disruption of the hydrogen bond. Among the top 100 modes with lowest free energy in each of Cys101-Ile152$_{A\&B}$ and Cys101-Ile152$_{B}$, we identified ten modes that showed least overlap between the Cys101-Ile152$_{A\&B}$ and Cys101-Ile152$_{B}$ (Extended Data Figure S6). We then computed root mean square fluctuations (RMSF) resulting from sampling these motion modes (Figure 4F). Interestingly, we observed that the perturbations in the hydrogen bonding network are propagated primarily to the IJ-linkers and helix J, consistent with the MD simulations. Strikingly, the largest RMSFs within the IJ linkers were observed near the active site in the opposite protomer, suggesting that the two sites are in allosteric communication. Identical analyses on the B-protomer hydrogen bond or the XFEL structure yielded similar results. By contrast, ten randomly selected motion modes lead to conformational changes distributed non-specifically throughout the dimer (Extended Data Figure S6).

Our results indicate that the conformational changes upon Cys101-Ile152 H-bond modification in the synchrotron structure correspond to later steps in the catalytic cycle, allowing helical motion that facilitates intermediate hydrolysis and product release. At the same time, increased allosteric transmission during these later steps may prime the dimer for the next catalytic cycle.

**Kinematic Flexibility Analysis (KFA) of the conformational ensembles modulated by the C101-I152 hydrogen bond.**

In KFA[4], a protein is represented as a kinematic linkage, with rotatable bonds $\mathbf{q}$ as degrees of freedom (DoFs) and hydrogen bonds and hydrophobic interactions as constraints. The constraints introduce cycles in the kinematic linkage, which impose coordinated motion on the degrees of freedom to maintain the constraints. These coordinated motions take place on a lower dimensional manifold of conformation space. Briefly, *m* constraints in a protein with *d* degrees of freedom define a constraint manifold

$$\mathcal{Q} = \{\mathbf{q} \in \mathbb{T}^d | \mathbf{\Phi}(\mathbf{q}) = \mathbf{0} \in \mathbb{R}^m\}$$

Formally differentiating with respect to time yields a linear relationship between instantaneous changes in the DoFs ($\dot{\mathbf{q}}$, ‘velocities’) and corresponding changes in the constraints:

$$\frac{\mathrm{d}\mathbf{\Phi}}{\mathrm{d}t} = \mathbf{J}\dot{\mathbf{q}}$$

The matrix $\mathbf{J}$ is known as the constraint Jacobian. By way of a singular value decomposition $\mathbf{JV} = \mathbf{U\Sigma}$, we obtain an expression $\mathbf{J}\mathbf{v}_i = \sigma_i \mathbf{u}_i$ relating a change in molecular conformation $\mathbf{v}_i$ to a change in the geometry $\mathbf{u}_i$ of all hydrogen bonds and hydrophobic constraints. Instantaneous changes of DoFs $\mathbf{v}_i$ that do not affect hydrogen bonds and hydrophobic constraints lie in the nullspace of $\mathbf{J}$: $\mathbf{J}\mathbf{v}_i = \mathbf{0}$. Normalized singular values $\sigma_i$ are proportional to the magnitude of internal hydrogen bond energy changes for motions along each mode $i$, with the constant of proportionality equal to 3.24 kcal/mol, independent of the mode. This leads to a formal, dimensionless expression for changes in the free energy $\Delta F_i = \sigma_i - c_T s_i$, for each motion mode. Here,

$$s_{v_i} = \frac{1}{d} \exp\left\{-\sum_{j=1}^{d} \kappa_{ij} \log(\kappa_{ij})\right\}$$

is the normalized exponential of the Shannon entropy calculated for each motion mode, and

$$\kappa_{ij} = \frac{v_{ij}^2}{\sum_{j=1}^{d} v_{ij}^2}$$

are the normalized velocities for each mode $i$, and $c_T = 1$ is a constant. The Shannon entropy $s_{v_i}$ represents how contributions of DoFs for each mode are distributed over the protein (*mode collectivity*) and renders $\Delta F_i \in [-1,1]$ for $c_T = 1$. Thus, $\Delta F_i$ balances enthalpic (constraint relaxation) contributions and conformational entropy (collectivity of motions) for each mode. For details, see [4].

We compared the spectrum of motion modes in the case where both protomers of the XFEL structure are subject to the Cys101-Ile152 hydrogen bond as a constraint, to the case where only the B protomer has the Cys101-Ile152 hydrogen bond as a constraint. This situation corresponds to an event of instantaneous disruption of the H-bond in one protomer. To investigate how motion modes changed upon disruption of the H-bond, we identified ten modes from a surface of near-constant free energy changes that are most dissimilar between the two scenarios. More precisely, we calculated the matrix product $\mathbf{D} = \mathbf{V}_{\boldsymbol{B}}^{\mathbf{T}}\mathbf{V}_{\boldsymbol{A\&B}}$, where $\mathbf{V}_{\boldsymbol{A\&B}}$ are the right singular vectors of the XFEL structure when the Cys101-Ile152 H-bond is present in both protomers, versus $\mathbf{V}_{\boldsymbol{B}}$ for the B protomer only. Note that $\mathbf{D}$ is orthonormal, and that an entry $|\mathrm{d}_{ij}|$ near 1 denotes that $\mathbf{v}_{B,i}$ and $\mathbf{v}_{A\&B,j}$ are motions along the same direction in conformation space, whereas $|\mathrm{d}_{ij}|$ near 0 denotes orthogonal motions (Extended Data Figure S6).

We then identified motion modes that significantly changed or shifted in the spectrum between **B** and **A&B**. Those motions will have a different effect on the geometry/energy $\boldsymbol{u}_i$ of all hydrogen bonds and hydrophobic constraints and are differentially accessible to the protein after disruption of the Cys101-Ile152 hydrogen bond. Modes in the nullspace of $\boldsymbol{J}$ (Extended Data Figure S6, mode number > ~1,450 corresponding to vanishing singular values) are least conserved between $\boldsymbol{V_B}$, and $\boldsymbol{V_{A\&B}}$, mostly because the nullspace basis is not unique. These modes are generally localized, with low collectivity and unfavorable free energy changes. Therefore, we first identified a set of 100 modes with lowest free energy, from which we then selected the top ten modes with least overlap to any other mode, i.e., for which $max_j\,|d_{ij}|$ is smallest. The free energy and mode overlap of these ten modes are indicated with stars in Extended Data Figure S6. We then perturbed the XFEL structure with one individual step along each of these ten modes $\boldsymbol{v}_i$ with unit step size to obtain an ensemble in which the root-mean-squared fluctuations represent the overall change in mobility due to the additional or missing hydrogen bond.

**Extended Discussion**

Cysteine-gated conformational changes in ICH alter the active site environment, likely promoting progress along the reaction coordinate. ICH catalyzes a reaction that can be divided into an early phase dominated by nucleophilic attack of Cys101 at the electrophilic carbenoid carbon of its isocyanide substrate and a later phase dominated by hydrolysis of the thioimidate intermediate to release the N-formamide product (Figure 1). The early phase requires a reactive cysteine residue to initiate nucleophilic attack, while the subsequent phase requires water attack at the thioimidate and weaker nucleophilicity of Cys101 (*i.e.* a better leaving group) to release the product. These two phases of ICH catalysis place conflicting demands on the physical properties of Cys101. The divergent pre-steady state kinetics of the G150A and G150T mutants suggest a model where the strained helical conformation of ICH has the highest competence for the initial isocyanide attack by Cys101, forming the thioimidate intermediate (Figure 1). This is also consistent with the diminished propensity of Cys101 for photooxidation in G150T, where the helix is constitutively shifted, suggesting that Cys101 is less reactive in this environment. After formation of the thioimidate, the helix samples the shifted conformation due to weakening of the Ile152-Cys101 H-bond, dynamically remodeling the ICH active site (Figure 4). Evidence for displaced helical conformations in ICH crystals with a thioimidate intermediate from the XFEL experiment indicates that the active site adopts conformations that favor water entry and hydrolysis of the thioimidate intermediate. Upon water attack at the thioimidate to form a tetrahedral intermediate, the H-bond between Cys101 and Ile152 that defines the strained conformation of helix H can reform, thereby stabilizing the nascent Cys101 thiolate and making it a better leaving group. With the strained helical conformation restored, the product is released and leaves the active site poised for another cycle of catalysis (Figure 4). The synchrotron X-ray crystallographic data indicate that the G150A mutation enhances sampling of shifted helical conformations even in the absence of Cys101 modification, and this shifted conformation is further populated once the Cys101-Ile152 H-bond is weakened by Cys101-SOH formation. In addition, G150A ICH accumulates a spectrally distinct 335 nm species that we propose corresponds to the thioimidate intermediate detected by XFEL crystallography. Our interpretation of these data is that the thioimidate intermediate accumulates in G150A ICH owing to an impaired resetting of the strained helical conformation, which reduces the rate of thioimidate hydrolysis and enzyme turnover. Consistent with the pre-steady state kinetic data, this kinetic model for G150A ICH predicts that the early chemical steps promoted by the strained conformation of helix H would not be impaired by the mutation but that the steady-state rate would be significantly diminished. In contrast, the G150T mutation causes a constitutively shifted helix, reducing rates of both initial Cys101 attack at the isocyanide carbon atom in the first chemical step and thioimidate hydrolysis in later steps, as evidenced by the lower burst and steady state rates of G150T ICH. Therefore, the G150A and G150T mutations have divergent effects on the early steps of ICH catalysis but similar detrimental effects on the later, rate-limiting steps.

Modulation of protein dynamics is a powerful way to regulate protein function, as has been characterized in various systems. To our knowledge, ICH is the first example of a non-disulfide cysteine modification regulating functional protein conformational dynamics. Nevertheless, conceptually similar examples of gated conformational dynamical changes exist. Redox-gated changes in flavoprotein structure and dynamics may play a major role in electron transfer by these proteins[5], and similar electron- or charge-coupled gating events occur in diverse systems[6-8]. Photoactivatable tags that modulate sampling of active enzyme conformations have been used to create catalytically enhanced enzymes[9]. In the DJ-1 superfamily to which ICH belongs, Cys106 oxidation to Cys106-$SO_2^-$ in DJ-1 results in little change in global protein conformation but stabilizes the protein by over 12 °C[10]. This stabilization is thought to be due to a strong (2.47 Å) hydrogen bond between Cys106-$SO_2^-$ and Glu18 that forms upon oxidation, reducing protein dynamics and stabilizing the protein.

More generally, transient covalent modification of proteins changes their potential energy surface. Therefore, various covalently modified species of a protein in the cell are likely to be dynamically distinct, providing another mechanism of diversifying protein function. The many potential covalent modifications of cysteine make this residue of particular importance for understanding how cellular signaling states, metabolite pools, and stress conditions couple to functional protein dynamics through the modification of amino acids in proteins. Future work on ICH and other cysteine-containing proteins will illuminate the diverse mechanisms by which cysteine-gated conformational changes can regulate protein function.

### References


1 Karplus, P. A. & Diederichs, K. Linking crystallographic model and data quality. *Science* **336**, 1030-1033 (2012).

2 van den Bedem, H., Bhabha, G., Yang, K., Wright, P. E. & Fraser, J. S. Automated identification of functional dynamic contact networks from X-ray crystallography. *Nature methods* **10**, 896-902 (2013).

3 Budday, D., Leyendecker, S. & van den Bedem, H. Geometric analysis characterizes molecular rigidity in generic and non-generic protein configurations. *J Mech Phys Solids* **83**, 36-47 (2015).

4 Budday, D., Leyendecker, S. & van den Bedem, H. Kinematic Flexibility Analysis: Hydrogen Bonding Patterns Impart a Spatial Hierarchy of Protein Motion. *J Chem Inf Model* **58**, 2108-2122 (2018).

5 Toogood, H. S., Leys, D. & Scrutton, N. S. Dynamics driving function: new insights from electron transferring flavoproteins and partner complexes. *FEBS J* **274**, 5481-5504 (2007).

6 Danyal, K., Mayweather, D., Dean, D. R., Seefeldt, L. C. & Hoffman, B. M. Conformational gating of electron transfer from the nitrogenase Fe protein to MoFe protein. *J Am Chem Soc* **132**, 6894-6895 (2010).

7 Catterall, W. A., Wisedchaisri, G. & Zheng, N. The chemical basis for electrical signaling. *Nat Chem Biol* **13**, 455-463 (2017).

8 Liu, Y. *et al.* A pH-gated conformational switch regulates the phosphatase activity of bifunctional HisKA-family histidine kinases. *Nat Commun* **8**, 2104 (2017).

9 Agarwal, P. K., Schultz, C., Kalivretenos, A., Ghosh, B. & Broedel, S. E. Engineering a Hyper-catalytic Enzyme by Photoactivated Conformation Modulation. *J Phys Chem Lett* **3** (2012).

10 Lin, J., Prahlad, J. & Wilson, M. A. Conservation of oxidative protein stabilization in an insect homologue of parkinsonism-associated protein DJ-1. *Biochemistry* **51**, 3799-3807 (2012).
